# HIV Virion Capturing Liposomes for Therapeutic Vaccination

**DOI:** 10.1101/2025.11.24.690259

**Authors:** Ted Keunsil Kang, Charles G. Ang, Gabriela Canziani, Heba Elkateb, Divine Thomas, Derek Yang, Amos B. Smith, Elias K. Haddad, Irwin Chaiken, Peter Deak

**Affiliations:** Drexel University, Department of Chemical and Biological Engineering, Philadelphia PA; Drexel University College of Medicine, Department of Biochemistry and Molecular Biology, Philadelphia PA; Drexel University, School of Biomedical Engineering, Philadelphia PA; University of Pennsylvania, Department of Chemistry, Philadelphia PA; Drexel University College of Medicine, Departments of Medicine and of Microbiology and Immunology, Philadelphia PA

## Abstract

HIV infection currently has no effective cures and requires lifelong antiretroviral treatments. Cures have failed due to HIV’s immune evasion and rapid mutation rate. Here we present a first in class HIV therapeutic vaccine, termed nanotrap therapeutic vaccines (NTVs) that are designed to capture circulating HIV virions and facilitate internalization by local antigen presenting cells. NTVs are modified liposomes that display the CD4 mimetic molecule, CJF-III-288, on their surfaces and have the TLR 7/8 agonist R848 loaded into their core. We show that NTVs can 1) bind gp120 and capture pseudoviral particles, 2) facilitate uptake by antigen presenting cells and 3) generate robust anti-HIV CD8 T cell immunity in transient infection mouse models. NTVs have translational potential to generate patient-specific HIV immunity.

## Introduction

Despite significant advancements in HIV antiretroviral therapies (ART) that render HIV highly treatable, true cures or preventative vaccines for HIV remain elusive, mainly due to HIV’s capacity for immune evasion and rapid mutation.^1^ Recent efforts have sought to leverage cutting edge immune-engineered preventative vaccine platforms to generate broad and hopefully effective HIV immunity.^2^ One recent example is Ad26.Mos4.HIV, an adenovirus type 26 vector delivering highly designed mosaic HIV antigens, which unfortunately demonstrated no efficacy in phase three clinical trials.^2^ Other efforts currently in clinical trials include nanoparticle protein formulations such as eOD-GT8 60mer and mRNA strategies, but each has yet to show clinical success.^3^

Alternatively, rather than a preventative vaccine, significant efforts have been made to generate a “therapeutic vaccine (TV),” i.e. a vaccine given after infection to stimulate immunity.^1^ This is also a tremendous challenge; HIV avoids immunity through a sophisticated combination of glycan shielding, mutagenesis and latency.^4–6^ Initial TV efforts, known as “shock and kill” sought to broadly stimulate immunity by administering strong immunogenic compounds, simultaneously releasing latent HIV reservoir and reactivating potent immunity, but this also failed in the clinic due to a combination of high toxicity and poor efficacy.^7,8^ More refined efforts have used combinations of broadly neutralizing antibodies(VRC07 and others) or T cell epitopes (HIVACAT T-cell immunogen) along with toll-like receptor 7 (TLR7) immunostimulants, with mixed efficacy in animal models.^9,10^ Alternatively, genetic editing strategies are being used to remove HIV from latently infected cells to generate a cure,^11^ but deploying these editing strategies selectively is a significant challenge, as HIV reservoir sites are highly anatomically diverse.^12^

While some of these efforts are still ongoing, current TV approaches still do not address the fundamental mutagenesis problem. Broadly reactive epitopes and antibodies have been identified, but ultimately lead to escape mutations.^13,14^ It is also unclear what is the “optimal” anti-HIV immunity that generates effective, suppressive responses to the virus that are sufficient to prevent spreading of infection and symptoms without ART. There is evidence that strong IgG responses can prevent infection; however, there is mounting evidence that strong CD8 T cell responses, rather than B cell responses, are critical to eliminate viral reservoirs and eliminate infected cells after infection has occurred.^15,16^ Elite controller (EC) patients, who control HIV infections naturally without ART, have both robust B and T cell responses, but there is evidence that anti-HIV CD8 T cell responses, driven by type I interferon (IFN-I) secreting dendritic cells (DCs), are responsible for effective viral control after initial infection.^17,18^ TV strategies to harness this “EC-like” immune response are currently underdeveloped.

Here we present a novel therapeutic vaccine candidate called nanotrap therapeutic vaccines (NTVs) that is designed as a strain agonistic TV that will skew HIV responses toward CD8 T cells. NTVs are modified liposomes multivalently displaying a small molecule CD4 mimetic, CJF-II-288, and loaded with a IFN-I skewing toll-like receptor (TLR) 7/8 agonist, R848.^19,20^ Rather than “shock and kill,” we propose designing novel nanoparticles to “bait and switch.” By decorating NTVs with molecules designed to mimic the main HIV entry receptor, CD4, we hypothesize that NTVs will capture circulating HIV by serving as “bait”. Then the HIV-NTV complex will be phagocytosed by local antigen presenting cells (APCs), leading to simultaneous presentation of relevant HIV epitopes and IFN-I-based activation of APCs, the “switch.” In this proof-of-concept study, we demonstrate that (1) NTVs can capture HIV viral particles, (2) facilitate specific uptake by DCs and (3) generate broad anti-HIV immunity in a transient infection mouse model. These data support the hypothesis that NTVs could generate functional strain-specific HIV immunity and validates that NTVs are a versatile and innovative first-in-class HIV therapeutic.

## Results

### Design of NTVs as modular platforms for viral capture

We developed a liposomal TV that could capture circulating HIV virions and facilitate the co-delivery of immune adjuvants and HIV virions to APCs to generate strain specific anti-HIV immunity. The NTV liposome consisted of three customizable components: 1) the HIV capturing moiety, 2) an immune agonist and 3) the liposome (**Fig. 1A-C**). We selected the HIV capturing moiety from a large body of literature of fusion inhibitors (i.e. molecules that bind envelope, env, proteins of HIV to prevent viral fusion).^21^ For our viral capture ligand, we chose CJF-III-288, a recently developed CD4 mimetic (CD4m) with both broad activity against a diverse set of HIV strains and high binding affinity (low nanomolar).^19^ For the immune agonist, we chose R848, a potent TLR7/8 agonist, known to stimulate IFN-I antiviral immunity and drive potent CD8 T cell responses in HIV.^22,23^ Clinically, however, excessive systemic IFN-I signaling is known to drive CD4 T cell depletion and lead to AIDS progression.^24^ To avoid this, R848 was loaded into liposomes, enhancing selectivity of immune activation to only cells that take up the virion.

**Figure 1.**
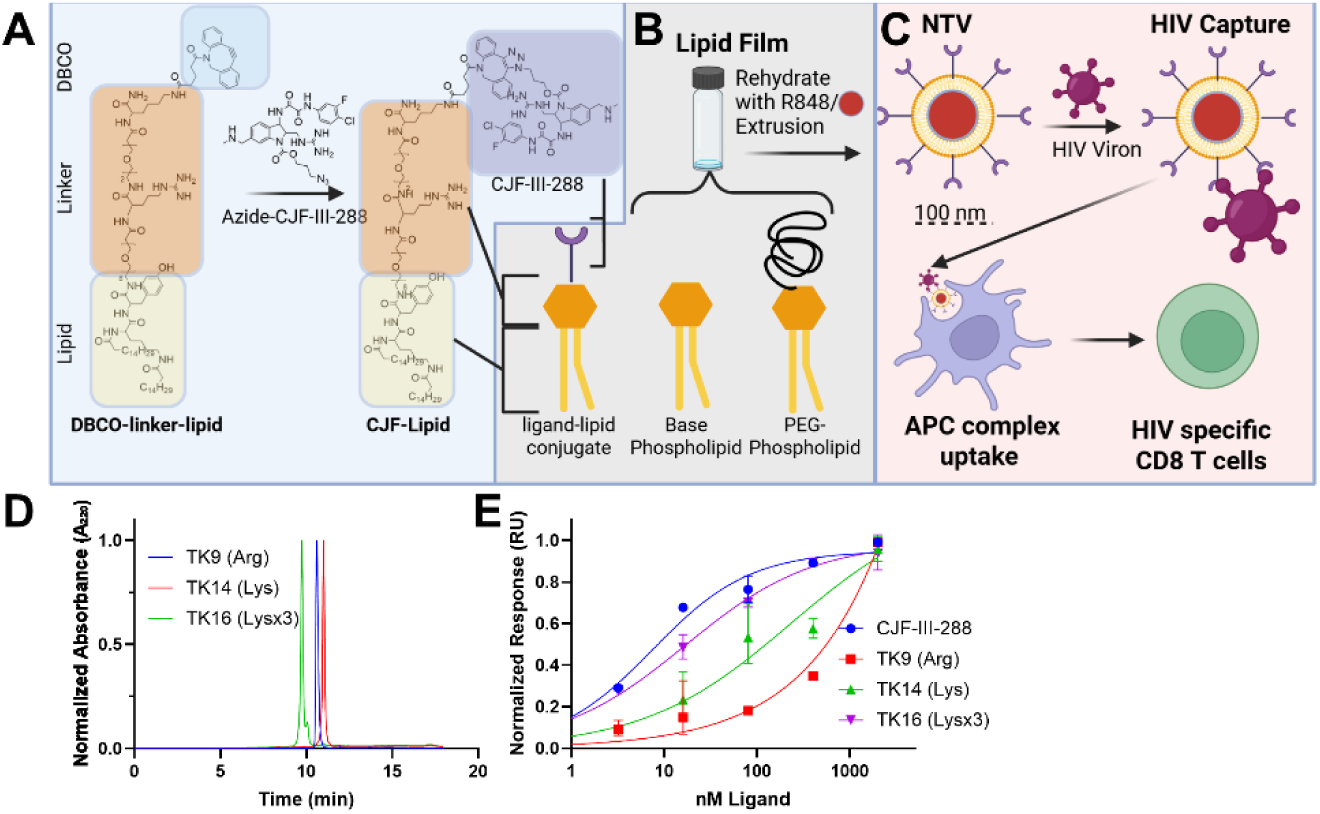
NTV Design/Proposed Mechanism and Chemical Analysis of CJF-lipid. (A) Schematic for synthesis of CJF-lipid conjugate, showing synthesis of DBCO conjugate including peptide/EG linker. Arg linker (TK-9) shown as example. (B) Design schematic of NTV synthesis. (C) Proposed NTV mechanism to generate anti-HIV immunity. (D) Representative HPLC Chromatograms at 280 nm for purified TK-9,14 and 16. (E) SPR analysis of CJF-lipids. 3000 RFU of gp120 YU2 variant was attached to a SPR chip (Biacore S200) and TK-16 (Lysx3 linker) binding was analyzed.

liposome complex.^25,26^ Furthermore, liposomes allowed for multivalent display of the viral capture domain, facilitating selectivity and specificity of viral capture.^27^ While this proof of concept study evaluated only one type of viral capture ligand (CJF-III-288) and one agonist (R848), it should be noted that NTVs are intentionally designed to be modular, and that all three components - ligand, agonist, and liposome platform - can be readily altered.

### Modified CJF-III-288 bind gp120

Our first goal in designing the NTV system was to generate modified CJF-III-288 molecules that display lipid tails to facilitate insertion into liposomes and proper surface display to capture virions (**Fig. 1A/B**). To this end, azide modified CJF-III-288 molecules (DY-IV-040) were synthesized, then reacted with DBCO displaying linker-lipids using solid phase peptide synthesis (**Fig. 1A, Fig. S-1).**^28^ The linker-lipids contained a branched palmitic acid (C16) tail to facilitate liposomal insertion, polyethylene glycol spacers (PEG8) and positively charged amino acids to facilitate proper ligand display.^29^ For our initial studies, we synthesized linkers bearing arginine (Arg, TK9), lysine (Lys, TK14) or three lysine (Lysx3, TK16), capped with a DBCO linker. CJF-lipids were synthesized via azide-DBCO click chemistry, purified by HPLC and validated with HPLC/ MS (**Fig. 1D, Fig. S-2/3/4).** CJF-lipids were further validated by surface plasmon resonance (SPR) for gp120 binding by measuring the disassociation constant (K_d_). We observed a decreased binding affinity for TK9 and TK14, but only a small reduction in binding from unmodified CJF-III-288 for TK16 (9 vs 18 nM K_d_, **Fig. 1E, Fig. S-5, Table 1**).

**Table 1:** Binding Affinities of CJF-lipids with YU-2 gp120.

| CJC-lipid | Linker | K<sub>d</sub> (nM) | 95% CI (nM) |
| --- | --- | --- | --- |
| CJC-III-288 | N/A | 9.11 | 6.3 to 13.0 |
| TK9 | Arg | 502 | 247 to 793 |
| TK14 | Lys | 105 | 54.0 to 205 |
| TK16 | Lysx3 | 17.5 | 13.4 to 22.8 |

### NTVs display CJF-III-288, bind gp120 and HIV pseudoviral particles

We next synthesized NTVs *via* the thin-film extrusion method and validated binding to gp120 with three different assays.^28^ All liposomes contained bulk DSPC (Distearoylphosphatidylcholine) loaded with 5% DSPE-mPEG2000, 5% cholesterol and 0.2% DiD dye. NTVs contained between 2-0.01% CJF-lipid molecules while blank liposomes did not. CJF-lipids loaded into NTVs and generated liposomes of approximately 140-160 nm hydrodynamic diameter when extruded through 100 nm pores as measured by dynamic light scattering (DLS, **Table 2, Fig. S-6/7**).

**Table 2:**
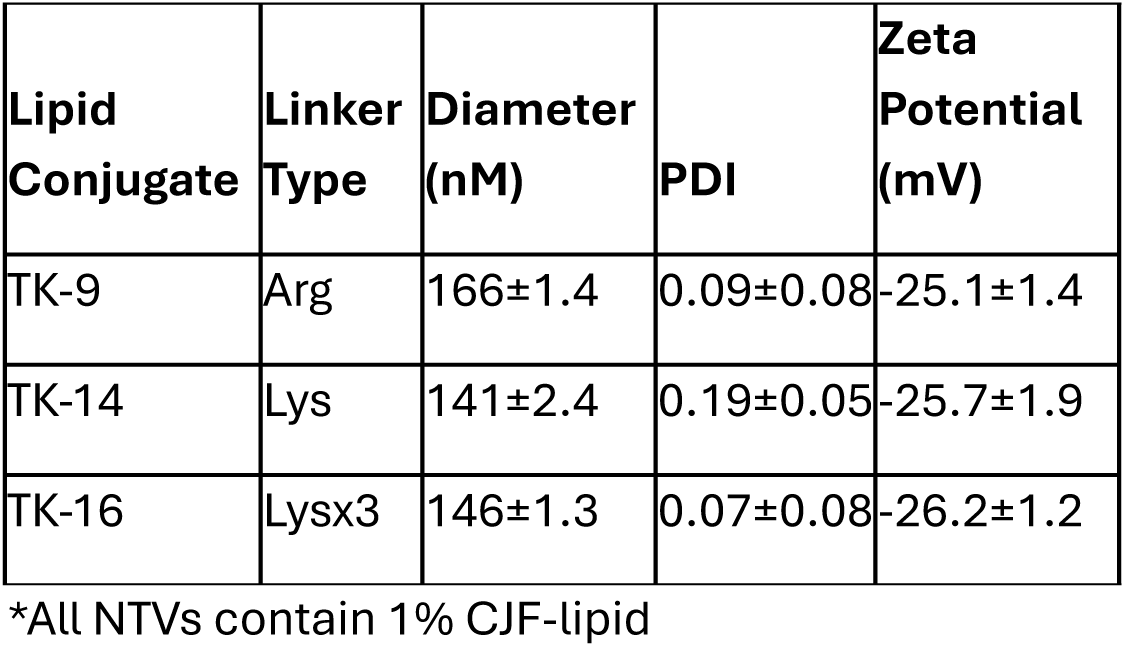
NTV Characterization.

Gp120 binding was analyzed first by incubating NTVs with TF228.1.16 (TF228), a stably transfected lymphoma cell line that expresses a full length gp160 (BH10 isolate) on its surface.^30^ We generated DiD labeled NTVs bearing 1% of TK9, TK14, TK16 or a control lipid (TK14 linker without CJF-III-288) or bearing no CJF-lipid (blank) and incubated them with TF228 cells at 37°C for 1 h, then analyzed with flow cytometry (**Fig. 2A/B**). We observed the highest uptake from TK9 and TK16 NTVs and little to no uptake from control liposomes. We generated NTVs varying TK16 loading ratios and repeated this experiment, observing increased binding as percentage loading (**Fig. 2C**). We identified 1% TK16 as the optimal NTV and unless noted we will use this formulation for the remainder of the study.

**Figure 2.**
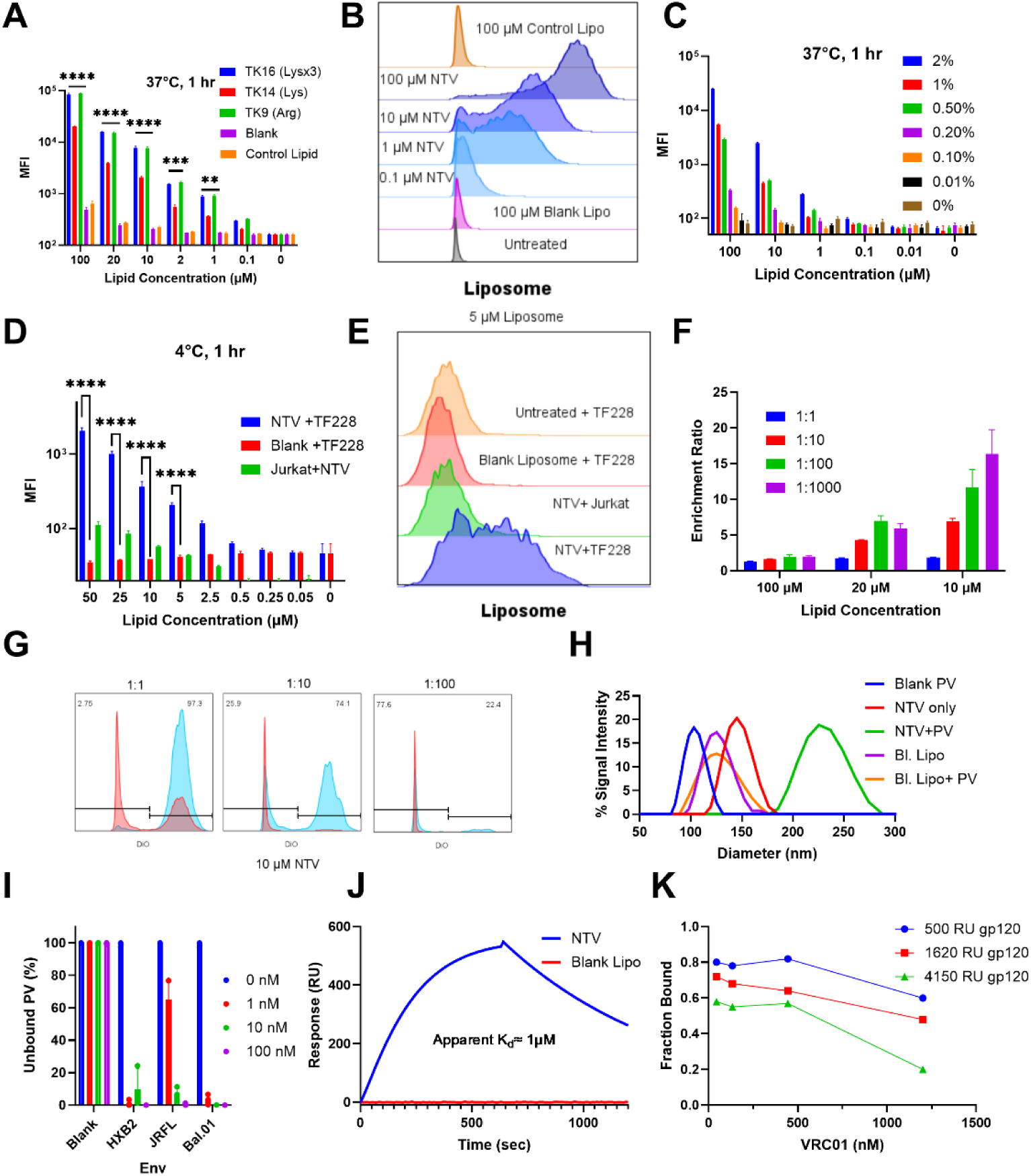
NTVs bind to gp120 and HIV pseudoviral particles. (A) 100k TF228 cells (bearing gp120, TF228.1.16) were incubated with DiD-labeled NTVs loaded with 1% CJF-lipids, a control lipid (TK14 without CJF moiety) or blank liposome for 1 h at 37°C. Uptake and binding were determined by flow cytometry. (B) Representative histograms of A for TK16 loaded NTVs. (C) Similar to part A, but NTVs were synthesized with 0.01-2% TK16 lipid. (D) 100k TF228 (gp120 bearing) or Jurkat (gp120 negative) were incubated with NTVs bearing 1% TK16 or blank liposome. (E) Representative histograms at 5 µM total lipid concentration. (F) NTVs selectively target gp120 expressing cells. DiO-labeled TF228 and Jurkat cells were co-cultured at 1:1, 1:10, 1:100, and 1:1000 ratios then treated with DiD-labeled 1% TK16, 100 nm NTVs for 1 h at 37°C. An enrichment ratio (%DiO+ of DiD+ cells / %DiO+ of all cells) was calculated. (G) Representative histograms of DiO signal from all cells (red) and DiO+ cells (blue). (H) SPR analysis of NTVs/Psuedovirus (PV) complex formation. Replication incompetent PV (pNL4-3 backbone expressing HXB2 env) was stained with DiO and incubated at 0.5 ng/mL p24 concentration with 1 nM total lipid of blank liposome or NTVs (1% TK16) for 30 min at 37°C then analyzed via DLS. Representative plot of three independent experiments. (I) DLS complex analysis like part H but varying PV env type on pNL4-3 backbone. Unbound PV= signal from 50-200 nm. N=3 (J) SPR analysis of NTVs. 25 µM total lipid of 1% TK16 NTV was analyzed on SPR chip loaded with 500 RU of YU-2 gp120 protein. (K) SPR occlusion assay. SPR chip was loaded with 500, 1620 or 4150 RU of YU-2 gp120 then treated with varying concentrations of VCR01 anti-gp120 antibody, then treated with 25 µM total lipid of 1% TK16 NTV. The fraction of maximal signal compared to no VCR01 antibody was calculated. Significance was determined by two-way ANOVA with Dunnett’s test for multiple comparisons. **p<0.01, ***p<1×10^−4^, ****p<1×10^−5^

To further control for any non-specific interactions, we incubated either TF228 or Jurkat (non-gp120 expressing controls) with NTVs at 4°C and observed significantly increased uptake with NTV-TF228 samples compared to controls (**Fig. 2D/E)**. To further analyze the capacity of NTVs to selectively identify gp120 expressing cells, we performed similar experiments as in **Fig. 1A** with co-cultures of DiO labeled TF288/ unlabeled Jurkat varying ratio from 1:1 to 1:1000 then used flow cytometry to identify DiD+, DiO+ cells compared to DiO-, DiD+. We observed that NTVs selectively targeted TF228 cells and calculated an “enrichment ratio,” i.e. %DiD+ of DiO+ divided by % DiO+ population, which represents the increased odds of NTV labeling over random chance (**Fig. 2F/G**). It should be noted that while NTVs did not inhibit viral infection using standard GHOST assays, NTVs did not interfere with infection inhibition by clinically deployed ART(**Fig. S-8).**^31^

We next observed complex formation between non-infectious pseudoviral particles (PV) using dynamic light scattering (DLS).^32^ We incubated PVs with pNL4-3 replication deficient backbone transfected with HXB2 env at 1, 10 or 100 nM total lipid concentration of either blank liposome or NTV and observed increases in complex size, suggesting a 1:1 virus-NTV complex at 1 nM (**Fig. 2H**).^33^ Complex sizes varied with incubation time and NTV concentration (**Fig. S-9).** We further validated that NTVs, but not blank liposomes, bound PV with HXB2, Bal.01 and JFRL env and did not bind env deficient PVs (**Fig. 2I, Fig. S-9**).

SPR analysis of protein binding to NTV (25 µM total lipid) yielded an apparent K_d_ of ∼1 µM based on total lipid concentration, and we did not observe binding from blank liposomes (**Fig. 2J, Fig. S-10)**. Given an estimated 1% ligand incorporation, ∼50% external orientation, and ∼40% loading efficiency, the effective ligand concentration corresponds to ∼2 nM.^29^ This value likely reflects multivalent avidity rather than true monovalent affinity, as rebinding and ligand clustering on the liposome surface can markedly enhance apparent binding strength.^34^ We further performed a competition/occlusion assay with VCR01, a broadly neutralizing antibody that targets the conserved CD4-binding site (CD4bs) on gp120 and thus competes with CJF-III-288.^35^ We observed binding inhibition of NTVs when SPR chips were preincubated with VCR01, indicating that NTV specifically binds gp120 via CJF-III-288 (**Fig. 2K**). These data, taken together, strongly supported our hypothesis that NTVs bound gp120 *via* CJF-III-288 and could effectively capture HIV virions.

### R848 Loaded NTVs are internalized and stimulate APCs

We then loaded NTVs with the TLR7/8 agonist, R848, and analyzed APC uptake and activation. NTVs were formulated with an initial R848 loading concentration of 166 µM, yielding a final encapsulated concentration of 60 µM (36% encapsulation efficiency) after dialysis at a total lipid concentration of 1 mM (drug-to-lipid molar ratio = 1:16.7, **Fig. S-11).** To test our hypothesis that NTVs facilitate viral internalization via complex formation, 100k murine bone marrow derived dendritic cells (BMDCs) were incubated with DiO labeled HXB2 PV and NTV or blank liposomes for 1 h at 37°C and uptake measured by flow cytometry (**Fig. 3A, B, Fig. S-12)**. Murine cells were chosen as they are not susceptible to infection, limiting confounder effects on observed viral uptake.^36^ We observed significant increases in DiO+, DiD+ cells for NTV compared to blank liposomes. We repeated this assay with a 24 h incubation and observed increases in release of the cytokines TNF-α and IL-6 and expression surface markers CD80 and MHCII, all of which indicate APC immune activation, for both NTVs and blank liposomes loaded with R848 and free R848 as expected (**Fig. 3C-F**).^37,38^ Interestingly, we observed that NTVs without R848, generated immune activation at 100 µM. We hypothesized this effect is due to HIV viral RNA triggering TLR activation.^39^

**Figure 3.**
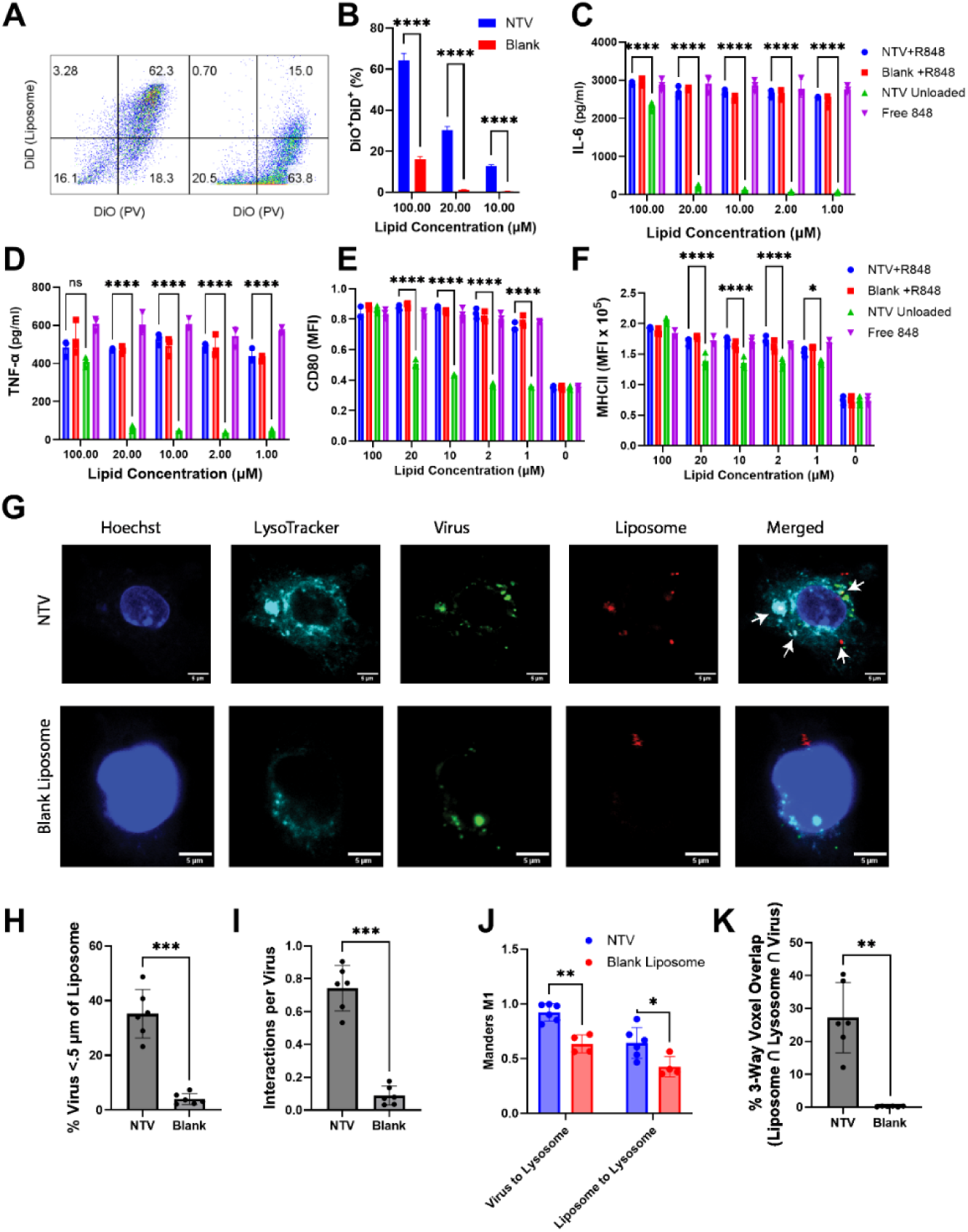
NTVs increase viral uptake and stimulate immune activation from antigen presenting cells. (A) DID labeled blank liposome loaded with R848, NTVs loaded with R848 or NTVs without R848 or free R848 at varying concentrations were incubated with 25 pg/mL p24 of DiO labeled HXB2 PV for 30 minutes then incubated with 10k BMDCs for 1 h, then analyzed with flow cytometry. Note R848 concentrations were selected to match total R848 delivered within liposomes at same lipid concentration. Representative flow plot is shown. (B) Quantification of %DiD+, DiO + cells. (C) Similar to part a, but instead BMDCs were incubated with virus-liposome complexes for 24 h, then supernatants isolated and IL-6 (D) TNFα, (E) and cells stained for the marker CD80 or (F) MHCII. (G) 100k BMDCs were attached to a glass coverslip and then incubated with the same liposome-virus mixture as in part A, but only at 20 µM total lipid. Cells were incubated for 1 h, washed, treated with lysotracker red per manufacturer’s instructions then washed and fixed with 2% PFA solution for 15 minutes, washed again and then imaged. Representative images of individual BMDCs are shown for NTV (top) and blank liposome (bottom) treated groups. White arrows in merged image indicate areas of co-association between lysosome, NTV and virus. (H-K) Quantification of 6 widefield confocal images of BMDCs per condition with each image containing 8-20 cells with 5-7 1 µm z slices. (H) Areas of liposome and virus were rendered as 3D objects and the % of virus within 500 nm of a liposome object was calculated. (I) Similarly, the average number of liposome associations per virus were calculated. (J) Using the same images, Manders M1 overlap coefficient was calculated for both virus-lysosome and liposome-lysosome. (K) The percentage of co-localization between lysosome, liposome, and virus voxel was calculated. Significance was determined by student t test for H, I, K or two-way ANOVA with Dunnett’s test for multiple comparisons for all others. *p<0.05, **p<0.01, ***p<1×10^−4^, ****p<1×10^−5^

To further interrogate our hypothesis that NTV-virus complexes are co-internalized, we performed confocal microscopy on BMDCs similarly treated as in **Fig. 3A** with DiO labeled PV and either NTV or blank liposome. BMDCs were stained with lysotracker staining dye (Thermo) and nuclear stain, fixed, imaged and analyzed with Fiji ImageJ to identify areas of overlap between virus, NTV and lysosome (**Fig. 3G, Fig. S-13**).^40^ To quantify overlap, we first generated a 3D distance map of virus and liposome signals. We then calculated significant increases for NTV compared to blank liposome in the percent of virus within 500 nm of liposome and the average number of liposomes interactions per virus (**Fig. 3H/I).** We also calculated a significant increase in co-localization of both virus in lysosome and liposome in lysosome by Mander’s coefficient (**Fig. 3J**).^41^ By analyzing voxel overlap between virus, liposome and lysosome signal, we calculated approximately 25% of all virus voxels had overlap with both liposome and lysosome signal for NTVs compared to less than 2% for blank liposome (**Fig. 3K**). It is also noted that NTV treated cells had higher average liposome signal and lower virus signal overall compared to blank liposomes, confirming our data in figure 2 for increased NTV uptake and suggesting that internalized virus may be rapidly degraded by lysosomes, whereas blank liposomes may demonstrate more non-specific membrane fusion **(Fig. S-14**). Taken together, these data supported our hypothesis that R848 laden NTVs drive complex formation with HIV PVs and were selectively internalized within lysosomal compartments *in vitro*.

### NTVs selectively target APCs in vivo and facilitate broad CD8 T cell immunity in transient infection mouse models

We performed two *in vivo* experiments to interrogate how NTVs target APCs and trigger immunity in vivo. For both *in vivo* experiments, C57Bl/6 mice were injected with replication deficient HXB2 env PV to simulate active early HIV infection and then injected with NTV 5 minutes later. While C57Bl/6 mice are not suspectable to HIV infection without modification of virus or animal, mice can generate effective anti-HIV immunity if given proper adjuvant co-stimulus.^42^ This “transient infection” model retained circulating HIV p24 for up to 4 h before clearance (**Fig. S-15)**. While this model did not fully recapitulate a full HIV infection, it was chosen as an experimentally simple test to validate if NTVs can facilitate anti-HIV immunity. To this end, we first injected mice with PV then NTV, free R848 or blank liposomes then monitored liposome uptake in RBC depleted blood at 1, 4 or 24 h post injection (**Fig. 4A/B)**. We then analyzed immune cell uptake in both circulating blood and spleen at 24 h (**Fig. 4C/D**). We observed increased uptake of NTVs compared to blank liposomes at 1, 4 and 24 h and an increased uptake specifically in circulating and spleen resident plasmacytoid DCs (pDC, CD11c+, B220+).^43^ It should also be noted that NTVs did not increase systemic inflammatory cytokines at 1 h post injection, alleviating toxicity concerns over injecting strong TLR agonists (**Fig. S-15**).^10^ This suggested that NTVs direct HIV virions towards uptake by pDCs, a cell type known to generate potent anti-viral CD8 T cell responses via IFN-I signaling.^44^

**Figure 4.**
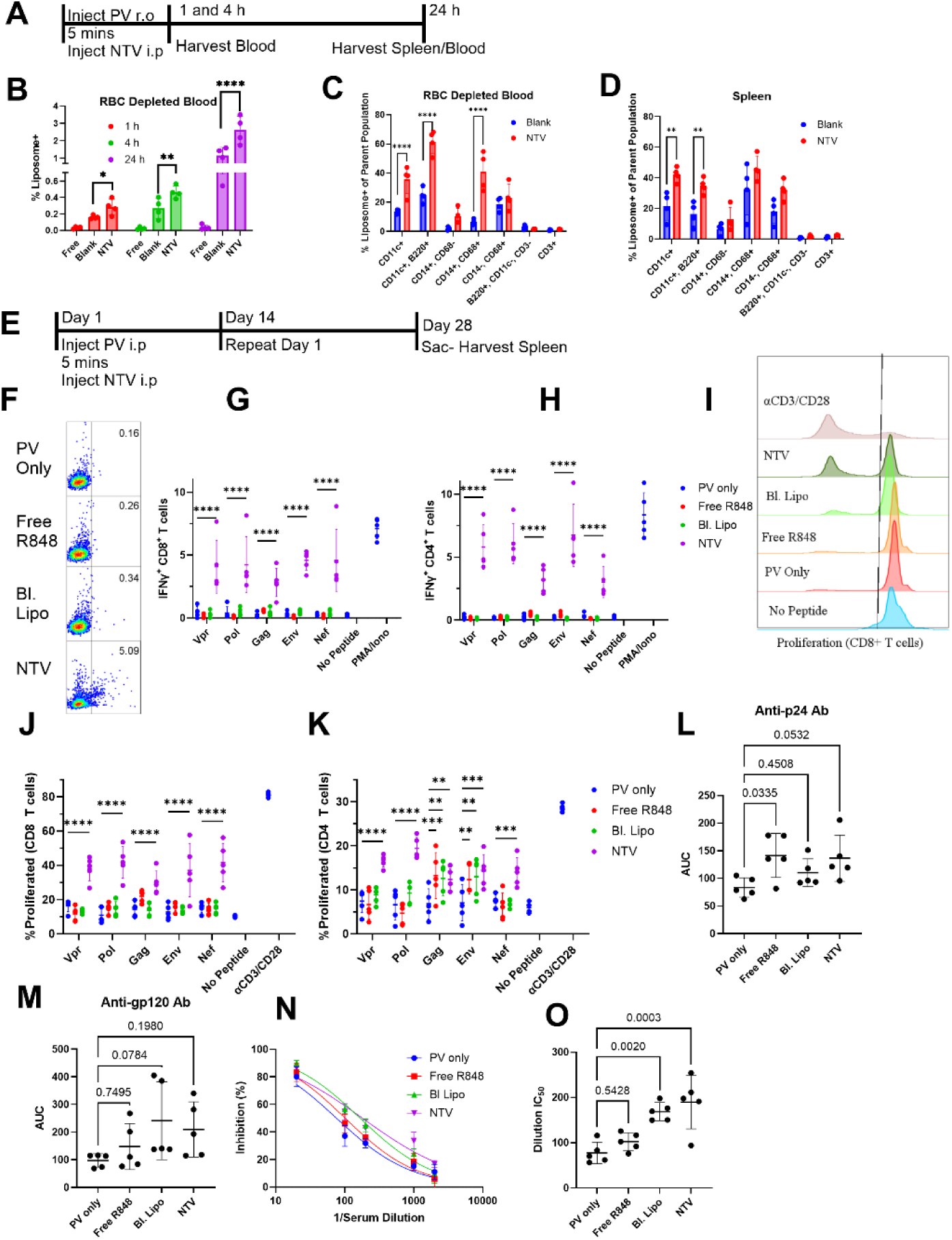
NTVs promote anti-HIV broad CD8+ T cell immunity in transient infection mouse models. (A) Schematic of in vivo uptake experiments. Briefly, C57Bl/6 mice (N=4) were injected r.o. with 0.1 ng p24 of BaL.01 PV, then injected i.p 5 minutes later 160 µMols total lipid of blank DSPC liposome loaded with 10 µMols of R848 (Blank) or similar amount of NTV (including 1% TK-16 total lipid). Blood was drawn via cheek bleed at 5 min, 1 h, and 4h post injection. At 24 h, mice were sacrificed, blood drawn and spleen isolated. (B) RBC depleted blood was analyzed via flow cytometry for liposomes. RBC Depleted blood (D) or disaggregated spleen cells (E) were analyzed for % liposome+ cells in the following cell populations: CD11c+, pDC (CD11c+, B220-), monocytes (CD11c-, CD68-, CD14+), CD14+ macrophages (CD14+, CD68+), CD14-macrophages (CD14-, CD68+), B cells (CD11c-, B220+) or T cells (CD3+). See Figure S-13A for gating strategy. (E) Schematic for in vivo vaccination experiment. C57Bl/6 (N=5) mice were injected i.p with 0.1 ng p24 of HXB2 PV on the left side. 5 minutes later, mice were injected i.p. on right side with either PBS (PV only), 160 µMols total lipid of blank DSPC liposome loaded with 10 µMols of R848 (Bl. Lipo) or similar amount of NTV (including 1% TK-16 total lipid). Additional control was a single injection of 10 µg of both p24 and gp120 protein from HXB2 mixed with 75 µMols R848 in PBS (Free R848). Both injections were repeated on day 14. Splenocytes were isolated and analyzed by (F-H) ICS and T cell proliferation (I-K). For ICS, cells were stimulated with peptide pools from 1 µg/mL Vpr, Pol, Gag, Env or Nef for 1 h, then blocked with brefeldin A for 5 h. Cells were fixed, stained and analyzed by flow cytometry for intracellular IFNγ. Controls include PMA/Ionomycin treated samples or cells treated with no peptide. (F) Representative flow plots of SSC (y axis) by IFNγ signal (x axis). (G) IFNγ+ CD8+ T cells, (H) IFNγ+ CD4+ T cells. For T cell proliferation, splenocytes were treated with same peptide pools plus 0.2 µg/mL anti-CD28. Positive controls were treated with 1 µg/mL of both anti-CD3/CD28 antibodies. Cells were stained with proliferation dye and incubated for 72 h, then analyzed by flow cytometry. (I) Representative flow plots of Live, CD8+ T cells with SSC (y axis) by Proliferation stain (x axis) (J) Proliferation+ CD8 T cells, (K) Proliferation+ CD4 T cells. (L) Serum isolated on day 28 was tested for Anti-p24 IgG or (M) anti-gp120 IgG and area under the curve (AUC) of serial dilutions calculated for each mouse. (N) Serum analyzed by GHOST cell inhibition assay and % inhibition calculated. (O) A dilution factor IC50 value for each mouse was calculated. Significance was determined by student t test for L, M, O or two-way

Finally, we sought to validate our hypothesis that NTVs would generate robust anti-HIV CD8 T cell immunity. To do this, we sequentially injected with HXB2 PV (100 pg p24) and then 160 µMols NTVs or blank liposomes (both containing 10 µMol R848) on day 1 and then on day 14 (**Fig. 4E**). An additional control of 10 µg of both p24 and gp120 protein from HXB2 PV mixed from 75 µMols of free R848 was also included (Free R848). After 28 days, mice were sacrificed and splenocytes analyzed via intracellular cytokine staining (ICS) and T cell proliferation after ex vivo stimulation with HIV peptide pools. ICS staining for IFNγ secreting T cells demonstrated significant increases in all peptide pool (Vpr, Pol, Gag, Env and Nef) simulated groups for both CD4 and CD8 T cells but not any other treatment group, suggesting NTVs trigger a potent anti-viral Th1 immune response (**Fig. 4F-H, Fig S-16A**).^45^ T cell proliferation data demonstrated a more diverse response however; whereas only CD8 T cells from NTV groups proliferated, CD4 T cell proliferation from both Free R848 and blank liposome groups significantly increased in response to Gag and Env simulation (**Fig. 4I-K**). Serum at day 28 was also analyzed for anti-p24 and gp120 IgG; only the free R848 group demonstrated significantly increased area under the curve (AUC) titer for p24 compared to PV only treated mice, although NTV did demonstrate a non-significant increase in both p24 and gp120 antibody titers (**Fig. 4L/M, Fig. S-16B/C)**. Serum antibodies were further analyzed with in vitro GHOST inhibition assays at various dilutions and a dilution IC_50_ value calculated for each mouse (**Fig. 4N/O, Fig. S-16D).**^46^ We observed a significant increase in the dilution IC_50_ value for both blank liposome and NTV treated groups, indicating more potently inhibitory antibodies. These results supported our hypothesis that NTVs generate broad CD8 T cell immunity, although interestingly showed less potent antibody responses.

## Discussion

This proof-of-concept study introduced a first-in-class TV candidate for HIV, NTVs. The data presented in this study demonstrate that NTVs can capture HIV PV particles, facilitate uptake into DCs and generate CD8 T cell skewed immunity. Particularly, it is this potent and broad CD8 T cell immunity that is uniquely promising for HIV TVs (**Fig. 4**). CD8 T cells are critical for HIV control and are a potential pathway for eliminating or reducing viral reservoirs in HIV patients.^47^ It has been noted in several studies that EC patients have increased IFN-I responses from pDCs leading to potent CD8 T cells that are critical for viral control.^48^ While NTVs cannot recapitulate the genetic components (i.e. HLA alleles etc.) responsible for these pDC responses in ECs, we hypothesize that NTVs can kick-start this pDC-IFN-CD8 T cell axis and potentially recapitulate the effective control of HIV responses. It should also be noted that the CD8 T cell response was very epitope diverse, spanning multiple important regions of HIV. This is encouraging for clinical transition, specifically for clearing latent reservoirs.^15^

A unique strength of NTVs that differentiates it from other TV candidates is utilizing circulating virions as antigen. Critically, this eliminates the need to predict antigens, but also the multivalent design allows for relatively small doses to remain effective. Due to limited availability of source materials, we were conservative with both our CJF-lipid loading (only 1% of total lipid) and total liposome doses. The apparent K_d_ of 1% loaded TK16 NTVs was approximately 2 nM, but due to multivalent effects for binding and rebinding, we observed formations of virus-NTV complexes as low as 1 nM total lipid (or 2 pM CJF-lipid once accounting for loading and external presentation, **Fig. 2**). Due to high binding, it is possible that NTVs could be used to identify low levels of circulating virus or even use the low env expression to identify recently reactivated viral reservoir cells.^12,49^ As we show in Figure 2F, NTVs are selectively uptaken by a nearly 20-fold ratio by env expressing cells when the ratio of env+ to env-cells is 1:1000. While for most ART controlled patients, the ratio of latent cells to uninfected is approximately 1:10^6^, highlighting the need for further optimization, NTVs represent a potential tool for identifying env expressing cells.^50^

It should also be noted that for our *in vivo* studies, NTVs used a very small amount of HIV antigen in the form of PV injection. We injected PV containing 100 pg of p24 antigen compared to the 10 µg of p24 antigen for the control group, a 100,000 times difference, and still observed similar p24 antibody titers and improved CD8 T cell responses (**Fig. 4**). For gp120, assuming a ratio of 50 p24 to gp120 monomers, we injected approximately 2 pg of gp120 protein; this difference increases to over 5 million times.^51^ Clinically, p24 serum concentrations range from approximately <0.1 pg/mL (ART suppressed) to 100 pg/mL for viremic HIV patients.^52^ In our mice, the serum concentrations were between 20-60 pg/mL, meaning that our models roughly approximated serum levels observed for untreated patient in the early stages of infection (**Fig. S-15B)**. Future work will evaluate if NTVs could effectively generate HIV immunity from lower circulating viral concentrations. A more realistic clinical vision for NTVs is as a first-line therapy to be co-administered with ART when viral titers are still high to generate anti-HIV immunity prior to robust establishment of latent reservoirs.^12^

The last unique strength of NTVs as a TV candidate is their versatility. Each major component of the NTV design can be readily altered and optimized to target HIV. For this initial study, we implemented a single gp120 targeting element at relatively low loading, but future work would deploy other HIV targeting molecules on the same NTV. This could allow NTVs to more selectively and specifically target HIV. Furthermore, the adjuvant can be readily replaced with other adjuvants or adjuvant combinations to “tune” the anti-HIV immune response, while keeping systemic toxicity low as these liposomes are directed selectively towards immune cells. We could also adapt the liposomal platform for other nanoparticle formats. Liposomes were chosen due to their ease of synthesis and compatibility for customization.^29^ Other similar platforms, such as lipid nanoparticles (LNPs), offer similar customization and can load different agonists more efficiently.^53^ Finally, it should be noted that NTVs could be applied to other viruses with known surface protein targeting molecules, such as hepatitis B/C, SARs-CoV-2 or herpes simplex.^54–56^

We must stress that this study is a preliminary evaluation of a new class of TVs and, as such, this study leaves several opportunities for further research. The transient infection mouse model in non-humanized mice does not model active HIV infection and cannot evaluate the capacity of NTVs to reduce reservoirs or control infection. To that end, we can make no conclusions about the efficacy of the anti-HIV immunity observed in Figure 4. This model, chosen due to logistical and practical considerations, is appropriate, however, to test our hypothesis that NTVs can generate anti-HIV immunity, and particularly CD8 T cell immunity, to HIV in vivo. Future studies will explore the efficacy of NTVs to control infection in humanized mouse models with infectious HIV.^57^ In a similar vein, we used BMDCs for our uptake analysis, which cannot become infected, and used only replication incompetent PV rather than live virus. This was also done for practical and safety reasons, but future studies will observe NTV binding to HIV virions taken from patient samples. Live virus from patients have a broader diversity of env strains, as well as variations in env surface concentrations as compared to PVs.^58^ Future studies should also evaluate more HIV stains, both lab strains and from primary samples, to further strengthen our claim that NTVs exhibit broad activity. Furthermore, mice and humans have slightly variant responses to R848, which can bind TLR8 and TLR7 in humans but only TLR7 in mice.^59^ Overall, these questions highlight the translational potential of NTVs and set a clear path for future studies.

By repurposing a broad spectrum CD4m molecule (CJF-III-288), we have shown that NTVs can capture HIV virus, facilitate the uptake of NTV-HIV complexes by APCs and generate anti-HIV CD8 T cell responses with simple in vivo models. This study is a first step in the evaluation of NTVs as a potential novel TV that can generate patient specific anti-HIV immunity.

## Materials and Methods

### Materials

Unless otherwise noted, all cell culture reagents and flow cytometry antibodies were obtained from Thermo Fischer Scientific. All chemistry reagents were obtained from Fischer Scientific. The p24 ELISA kits were purchased from Abcam. The DiO labeling kit was purchased from Thermo Fischer Scientific. The following reagent was obtained through the NIH HIV Reagent Program, Division of AIDS, NIAID, NIH: 1) TF228.1.16, ARP-11481, contributed by Drs. Zdenka Jonak and Steve Trulli, 2) Jurkat (E6-1) Cells, ARP-177, 3) Human Immunodeficiency Virus Type 1 (HIV-1) NL4-3 ΔEnv Vpr Luciferase Reporter Vector (pNL4-3.Luc.R-E-), 4) HRP-3418, Human Immunodeficiency Virus 1 (HIV-1) BaL.01 Env Expression Vector, ARP-11445, contributed by Dr. John Mascola, 5) Ghost (3) CCR3+ CXCR4+ CCR5+ Cells, ARP-3943, 6) Peptide Pool, Human Immunodeficiency Virus Type 1 Subtype B (Consensus) Vpr Region, ARP-12860, contributed by DAIDS/NIAID, 7) Peptide Pool, Human Immunodeficiency Virus Type 1 Group M (Consensus) Pol Protein, ARP-13087, contributed by DAIDS/NIAID, 8) The following reagent was obtained through BEI Resources, NIAID, NIH: Human Immunodeficiency Virus Type 1 (HIV-1) HXB2 p24 Recombinant Protein, ARP-13126, 9) Peptide Pool, Human Immunodeficiency Virus Type 1 Subtype B (Consensus) Gag Region, HRP-12425, 10) Peptide Pool, Human Immunodeficiency Virus Type 1 Subtype B (Consensus) Envelope Region, HRP-12540, 11) Peptide Pool, Human Immunodeficiency Virus Type 1 Subtype B (Consensus) Nef Region, HRP-12545, 12) HEK-293 Non-Transfected Cell Line, NR-9313, 13) Monoclonal Antibody IgG YZ23, ARP-12047.

### Synthesis of CJF-lipids

CJF-lipids were synthesized using Fmoc-based solid-phase peptide synthesis (SPPS) on rink amide resin (Sigma-Aldrich). After assembly of the peptide backbone containing PEG spacers and a palmitic acid lipid tail, the side-chain ivDde protecting group on the terminal lysine (head-facing residue) was selectively removed. The exposed lysine amine was then coupled to DBCO–TFP ester to introduce a HIV binding CJF molecule. The azide-bearing CD4-mimetic CJF-III-288 was subsequently conjugated via copper-free click chemistry (SPAAC), yielding the final CJF-lipid construct. Crude products were purified by reverse-phase HPLC (Agilent) and confirmed by MALDI-TOF (Bruker) mass spectrometry. See materials and methods supplemental for more information.

### NTV Synthesis

NTVs were prepared by thin-film hydration using DSPC (89%), cholesterol (5%), and mPEG-2000-DSPE (5%), with CJF-lipids incorporated at 1 mol% and R848 encapsulated within the aqueous core. The dried lipid films were hydrated in PBS at 66 °C and extruded through 100-nm polycarbonate membranes (cytiva) to generate uniform nanoparticles. Unencapsulated R848 and free CJF-lipids were removed by dialysis using a 10 kDa cutoff membrane (Thremo). Final NTV formulations were characterized by DLS (Brookhaven Instruments) for hydrodynamic size and PDI, and by HPLC (Agilent) to quantify R848 loading efficiency.

#### Pseudoviral Production

Plasmid Production. HIV backbone (pNL4-3.Luc.R-E-, HRP-3418; obtained from BEI Resources) and env (BaL.01, ARP-11445; obtained from BEI Resources and HXB2, HXB2 and JFRL wild-type Env, a kind gift from Dr. Joseph Sodroski) plasmids were amplified using standard bacterial propagation. Chemically competent *E. coli* were transformed with each plasmid and plated onto LB-agar plates, followed by incubation at 37 °C overnight. Single colonies were inoculated into 3 mL TB broth supplemented with 100 µg mL⁻¹ ampicillin and shaken at 250 rpm at 37 °C overnight. The resulting starter cultures were transferred into 500 mL TB broth (100 µg mL⁻¹ ampicillin) and shaken at 250 rpm at 37 °C overnight. Bacterial pellets were collected by centrifugation at 5,000 × g for 10 min at room temperature. Plasmid DNA was extracted and purified using the PureYield Plasmid Maxiprep System (Promega) according to the manufacturer’s protocol.

Pseudoviral Production. HEK293T cells (NR-9313; obtained from BEI Resources) were seeded at a density of 2.5 × 10⁵ cells mL⁻¹ in a T75 flask and incubated overnight to allow adherence. Cells were co-transfected with 8 µg of HIV backbone plasmid (NL4-3) and 4 µg of env plasmid (BaL.01, JFRL, or HXB2). Supernatants containing pseudotyped HIV particles were collected 48 h post-transfection, cleared by centrifugation at 1,200 × g for 8 min, aliquoted, and flash-frozen in liquid nitrogen. Viral preparations were stored at −80 °C until use.

### Surface Plasmon Resonance (SPR)

The kinetic interaction assay was performed using an SPR biosensor, Biacore 3000 (Biacore, Uppsala, Sweden) as described previously.^1^ The SPR experiments were conducted at 25 °C in 1X PBS buffer (1 mM KH2PO4, 10 mM Na2HPO4, 137 mM NaCl, and 2.7 mM KCl, pH 7.4) with 0.005% Tween 20. Initially, CM5 sensor chips were activated using a standard amine coupling method using 1-ethyl-3-(3-(dimethylamino)propyl) carbodiimide (EDC) and N-hydroxysuccinimide (NHS) solution. Briefly, carboxyl groups on the sensor surface were activated by injection of 50 µL of a solution of 0.2 M EDC and 0.05 M NHS at a flow rate of 5 µL/min. Next, the protein ligand gp120, YU-2, was diluted in low-pH solvent (10 mM sodium acetate pH 5.0) and passed over the activated CM5 chip surface for covalent capture for the desired number of response units. Post-gp120 capture, the chip was deactivated using 1 M ethanolamine, pH 8.5. The real-time interaction was measured by injecting CJF-lipids or NTV over these surfaces. Surface regeneration was achieved by two 10 μL injections of 15 mM HCl solution at a flow rate of 100 μL/min after the dissociation phase. All analyses were performed in triplicate or duplicate. The Sensorgram for direct binding was analyzed using BIA Evaluation v4.0 software provided by Biacore (GE Healthcare). Prior to calculation, the binding data were corrected for non-specific interaction by subtracting the reference surface data from the reaction surface data and were further corrected for buffer effects by subtracting the signals due to buffer injections from those due to sample injections. Individual kinetic parameters were obtained from three independent experiments.

### Dynamic Light Scattering (DLS)

PV samples, 250 µL of approximately 1 ng/mL of p24 were stained with 0.5 µL of DiO Stain solution (Thermo) for 5 minutes at room temperature, then washed with a 40 kDa Zebra desalting column (Thermo). PV were then centrifuged at 10,000g for 2 minutes to remove any large complexes. PV was then incubated with liposomes for 5-30 minutes at room temperature and analyzed. Particle size distributions were measured by photon cross-correlation spectroscopy using a NANOPHOX (Sympatec GmbH, Clausthal-Zellerfeld, Germany; model NANOPHOX, software PAQXOS 6.2.2). Samples were diluted filter sterilized PBS, pH 7.4 to a count-rate within the instrument’s recommended range (50–300 kcps). Measurements were performed in polystyrene cuvettes with a path length of 45 mm at 25.0°C with active temperature control. The dispersant refractive index n= 1.331 and viscosity 0.890 mPa·s and the particle refractive index n = 1.52 + 0.001i were entered into the software. For each sample, the instrument was configured to automatically repeat sub-runs until the vendor-defined “Normal” standard-deviation stop condition was met. Results represent three independent experiments.

#### TF228 Cell Culture Assays

Cell Culture. TF228.1.16 (TF228) and Jurkat T cells were cultured at 37 °C and 5% CO₂. TF228 cells were maintained in DMEM (CORNING) supplemented with 10% FBS (Sigma-Aldrich), while Jurkat cells were maintained in RPMI-1640 (Life Technology) supplemented with 10% FBS. Cultures were split every 2–3 days to maintain logarithmic growth.

gp120 Binding Assay. TF228 cells were plated at a density of 1 × 10⁶ cells mL⁻¹ in 96-well flat-bottom plates and incubated overnight to allow stabilization. DiD-labeled blank liposomes or NTVs (1 mol% TK-16) were pre-incubated with DiO-labeled HXB2 pseudovirus for 30 min at 37 °C and 5% CO₂ and then added to the cells. After 1 h of incubation at 37 °C and 5% CO₂, cells were washed twice with FACS buffer (PBS containing 1% BSA and 5% FBS) and analyzed by flow cytometry.

gp120 Specificity Assay. TF228 and Jurkat cells were plated at 1 × 10⁶ cells mL⁻¹ in 96-well flat-bottom plates and incubated overnight. DiD-labeled blank liposomes or NTVs (1 mol% TK-16) were pre-incubated with DiO-labeled HXB2 pseudovirus for 30 min at 37 °C and 5% CO₂ and added to the cells. The plates were incubated for 1 h at 4 °C to restrict endocytosis, washed twice with FACS buffer, and analyzed by flow cytometry.

#### GHOST infection Inhibition Assay

Cell Culture. GHOST (3) CCR3+ CXCR4+ CCR5+ Cells CD4⁺ cells were maintained at 37 °C and 5% CO₂ in DMEM (CORNING) supplemented with 10% FBS (Sigma-Aldrich), 100 µg mL⁻¹ G418, 100 µg mL⁻¹ hygromycin B, and 10 µg mL⁻¹ puromycin. Cells were passaged every 2–3 days to maintain logarithmic growth and detached using Trypsin-EDTA prior to splitting.

Infection Inhibition Assay. GHOST cells were seeded at 1 × 10⁶ cells mL⁻¹ in 96-well flat-bottom plates and incubated overnight to allow adherence. HIV inhibitors or DiD-labeled NTVs (1 mol% TK-16) were pre-incubated with BaL.01 pseudovirus for 30 min at 37 °C prior to addition to the cells. After 1 h of incubation at 37 °C and 5% CO₂, cells were washed twice with FACS buffer (PBS containing 1% BSA and 5% FBS) and analyzed by flow cytometry to quantify inhibition of pseudoviral infection.

#### BMDC Culture Assay

BMDC Culture. Bone marrow was harvested from 6weeks-old C57BL/6 mice and differentiated into dendritic cells (BMDCs) using supplemented culture medium: RPMI 1640 (Life Technologies), 10% HI-FBS (Sigma-Aldrich), Recombinant Mouse GM-CSF (carrier-free) (20ng/ mL; BioLegend), 2mM l-glutamine (Life Technologies), 1% antibiotic-antimycotic (Life Technologies), and 50μM β-mercaptoethanol (Sigma-Aldrich). Bone marrow cells were treated with ACK lysis buffer for 10min, washed, and plated with differentiation media at 1 million cells in 10mL culture media. Cells were used for experiments on days 5–9.

BMDC Uptake Assay. Bone marrow–derived dendritic cells (BMDCs) were seeded at 1 × 10⁶ cells mL⁻¹ in 96-well flat-bottom plates and incubated overnight to allow adherence. DiD-labeled blank liposomes or NTVs (1 mol% TK-16) were pre-incubated with DiO-labeled HXB2 pseudovirus for 30 min at 37 °C and 5% CO₂ and subsequently added to the cells. After 1 h of incubation at 37 °C and 5% CO₂, cells were washed twice with FACS buffer (PBS containing 1% BSA and 5% FBS) and analyzed by flow cytometry.

BMDC Upregulation Assay. BMDC Uptake. Bone marrow–derived dendritic cells (BMDCs) were seeded at 1 × 10⁶ cells mL⁻¹ in 96-well flat-bottom plates and incubated overnight to allow adherence. DiD-labeled R848-loaded blank liposomes, unloaded NTVs, R848-loaded NTVs (1 mol% TK-16), or free R848 were pre-incubated with DiO-labeled HXB2 pseudovirus for 30 min at 37 °C and 5% CO₂ before being added to the cells. After 24 h of incubation under the same conditions, BMDCs were washed twice with FACS buffer (PBS containing 1% BSA and 5% FBS) and analyzed by flow cytometry to assess CD80 and MHC-II upregulation.

BMDC Cytokine Production Assay. BMDC Cytokine Production Assay. Supernatants from the BMDC upregulation assay were collected after incubation. TNF-α and IL-6 (Thermo) concentrations were quantified using commercial ELISA kits (Thermo) according to the manufacturer’s instructions to evaluate BMDC activation induced by NTVs.

### Confocal Microscopy

Bone marrow–derived dendritic cells (BMDCs) or DC-THP-1 were plated at a density of 1 × 10⁵ cells mL⁻¹ on glass coverslips and incubated overnight for adherence. Cells were treated with 10 µM total lipid of DiD-labeled blank liposomes or nanotrap vesicles (NTVs; 1 % TK-16) that had been pre-incubated for 30 min with 0.5 ng mL⁻¹ p24 of DiO-labeled HXB2 pseudovirus prior to addition. Following a 2 h uptake period at 37 °C, cells were stained with LysoTracker Red (Invitrogen) according to the manufacturer’s instructions, fixed in 1.5 % paraformaldehyde for 15 min at room temperature, and washed in PBS. Imaging was performed on a Leica Stellaris 5 laser-scanning confocal microscope using a 63× oil-immersion objective (NA 1.4). For each condition, 8–25 cells were analyzed from 4–6 image stacks, each containing 4–5 z-slices acquired at 1 µm intervals under identical laser power and detector gain settings.

Average Intensity per Cell: Z-stacks were processed in Fiji/ImageJ (v1.54). Images were converted to 8-bit format and calibrated to 0.089 µm × 0.089 µm × 1 µm voxel dimensions. Background fluorescence was subtracted using a 100-pixel rolling-ball radius, and brightness/contrast were normalized across replicates. Mean fluorescence intensity was averaged across all z-slices and normalized to the number of nuclei per field, yielding the average viral or liposomal signal per cell.

Manders Coefficient: Manders’ overlap coefficients (M₁, M₂) were computed after light Gaussian smoothing (σ = 0.5 µm) using the *Coloc2* plugin in Fiji. Thresholds were automatically set by the Costes bisection algorithm.^2^ M₁ values represent the fraction of viral or liposomal signal coincident with lysosomes, providing an unbiased measure of 2D colocalization.

3D Distance Measurements: Binary masks were generated for each channel by automatic thresholding followed by morphological dilation to approximate object boundaries. Three-dimensional Euclidean distance mapping was conducted using the *3D Distance Map* suite in Fiji. Viral–liposomal proximity was defined as particles within 0.5 µm of one another. Quantifications included the percentage of viral puncta within 500 nm of liposomes and the mean number of liposomes within 500 nm of each virus. This analysis followed established methods for 3D spatial colocalization and proximity mapping.^3^

Voxel-Wise Volume Overlap: Voxel-based overlap was determined from segmented binary masks of viral, liposomal, and lysosomal signals. The percent overlap was expressed as

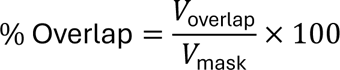

and triple-overlap (virus ∩ liposome ∩ lysosome) was computed by logical intersection of all three masks. This approach quantifies shared volumetric occupancy within the 3D cellular environment.^2,3^

### In Vivo biodistribution Assay

C57Bl/6 mice (N=4) were injected retro-orbitally (r.o.) with 0.1 ng p24 protein of Bal.01 env pseudoviral particles (BEI ARP-1144 env with BEI HRP-3418 backbone) in 100 µL. Mice were injected i.p 5 minutes later with 100 µL of 160 µMols total lipid of DiD labeled (0.2%) DSPC liposome loaded with 10 µMols of R848 (Blank) or similar amount of NTV (including 1% TK-16 total lipid) or 10 µMols of R848 free in PBS. Blood was drawn via cheek bleed at 5 min, 1 h, and 4h post injection. Blood was treated with 1x RBC lysis buffer (Thermo) and analyzed for DiD+ cells and cytokines using a LegendPlex mouse Th1/2 CBA kit (BD Biosciences). Mice were sacrificed at 24 h, spleens isolated, disaggregated with 100 mg/mL collagenase D and 20 U/mL of DNAse I in RPMI for 45 minutes, treated with 10 mM EDTA solution and then strained into a single cell suspension, and treated with RBC lysis buffer. Cells were then stained for CD11c+, pDC (CD11c+, B220-), monocytes (CD11c-, CD68-, CD14+), CD14+ macrophages (CD14+, CD68+), CD14-macrophages (CD14-, CD68+), B cells (CD11c-, B220+) or T cells (CD3+) and analyzed by Attune NxT flow cytometer (7 channel).

### In vivo vaccination assay

C57Bl/6 mice (N=5) were injected i.p on the left side with 0.1 ng p24 protein of HXB2 env pseudoviral particles (BEI ARP-1144 env with BEI HRP-3418 backbone) in 100 µL. Mice were injected i.p 5 minutes later with 100 µL of 160 µMols total lipid of DiD labeled (0.2%) DSPC liposome loaded with 10 µMols of R848 (Blank) or similar amount of NTV (including 1% TK-16 total lipid) or 10 µMols of R848 free in PBS mixed with 10 µg of both p24 and pg120 protein from HXB2. Mice were injected in this fashion on day 1 and day 14. Then on day 28, mice were sacrificed, blood drawn by cardiac puncture, spleens similarly isolated and processed into a single cell suspension as in biodistribution assay. Splenocytes were plated in 96 well flat bottom plates at 10^6^ cells per well in 200 µL of T cell media (RPMI 1640 supplemented with 10% heat-inactivated fetal calf serum (FCS/FBS), L-glutamine, and penicillin/streptomycin with 2 mM EDTA and 50 µM BME).

Intracellular Cytokine Staining (ICS): Splenocytes from the in vivo vaccination assay were treated with 1 µg/mL per peptide of HIV peptide pools (ARP12860, ARP-13087, ARP-13126, ARP-12425, ARP-12540, ARP-12545; obtained from BEI Resources) or 1x PMA/ionomycin solution (thermos) for 1 h, then treated with 1x eBioscience™ Protein Transport Inhibitor Cocktail for 4 h. Cells were then washed, stained with live/dead stain, fixed with 2% PFA for 15 minutes and stained for surface markers. Cells were permeabilized with 1x eBioscience™ Fixation/Permeabilization solution, then stained for IFNγ for 30 mins at room temperature, washed and analyzed via Attune NxT flow cytometer (7 channel).

T cell proliferation assays: Splenocytes from in vivo vaccination assay were stained with cell proliferation stain eflour 670 (Thermo) according to manufacturer’s instructions and then incubated with 0.2 µg/mL anti-CD28 antibody plus 1 µg/mL per peptide of HIV peptide pools. Positive controls were treated with 1 µg/mL of both anti-CD3/CD28 antibodies. Splenocytes were incubated for 96 h, washed and stained for surface markers CD69, CD3, CD4 and CD8, then stained for live dead stain and analyzed via Attune NxT flow cytometer (7 channel).

### Serum ELISA

Serum samples from the transient infection mouse model were collected on day 28 after centrifugation at 10,000 × g for 10 min. Anti-p24 and anti-gp120 IgG (ARP-12047; obtained from BEI resources) levels were quantified using commercial ELISA kits (Thermo) according to the manufacturer’s instructions. Antibody titers were calculated as the area under the curve (AUC) and compared with those of PV-only treated mice to evaluate the enhancement of humoral responses induced by NTVs.

### Serum GHOST Assay

GHOST cells were seeded at 1 × 10⁶ cells mL⁻¹ in 96-well flat-bottom plates and incubated overnight to allow adherence. Serum samples were pre-incubated with HXB2 pseudovirus for 30 min at 37 °C prior to addition to the cells. After 1 h of incubation at 37 °C and 5% CO₂, cells were washed twice with FACS buffer (PBS containing 1% BSA and 5% FBS) and analyzed by flow cytometry to quantify inhibition of pseudoviral infection.

## Acknowledgements

We thank the BEI program for facilitating access to standardized viral and immunological materials. We thank the Penn Chemistry: Mass Spectrometry Facility for help analyzing CJF-lipids and the Drexel College of Medicine SPR Shared Resource Facility for help with SPR analysis.

Funding for this project was provided by the NIH (NIAID) 1R56AI179226-01 to PD, IC and EKH.

## Author contributions

Conceptualization: PD, EKH, IC

Methodology: PD, IC, CA, DY, ABS, TKK

Investigation: PD, CA, TKK, HE, DT, GC, DY

Visualization: PD, TKK, GC

Supervision: PD, IC, EKH

Writing—original draft: PD, TK

Writing—review & editing: PD, IC, EKH

## Competing interests

All other authors declare they have no competing interests.

## Data and materials availability

All data are available in the main text or the supplementary materials.

## Notes

### Competing Interest Statement

The authors have declared no competing interest.

## References

1. Chen, Z. & Julg, B. Therapeutic Vaccines for the Treatment of HIV. Transl Res 223, 61–75 (2020).

2. Adepoju, V. A. et al. Navigating the Complexities of HIV Vaccine Development: Lessons from the Mosaico Trial and Next-Generation Development Strategies. Vaccines (Basel) 13, 274 (2025).

3. Cohen, K. W. et al. A first-in-human germline-targeting HIV nanoparticle vaccine induced broad and publicly targeted helper T cell responses. Sci Transl Med 15, eadf3309 (2023).

4. Chavez, L., Calvanese, V. & Verdin, E. HIV Latency Is Established Directly and Early in Both Resting and Activated Primary CD4 T Cells. PLOS Pathogens 11, e1004955 (2015).

5. Parajuli, B. et al. Identification of a glycan cluster in gp120 essential for irreversible HIV-1 lytic inactivation by a lectin-based recombinantly engineered protein conjugate. Biochem J 477, 4263–4280 (2020).

6. Abram, M. E. et al. Mutations in HIV-1 Reverse Transcriptase Affect the Errors Made in a Single Cycle of Viral Replication. J Virol 88, 7589–7601 (2014).

7. Kim, Y., Anderson, J. L. & Lewin, S. R. Getting the “kill” into “shock and kill”: strategies to eliminate latent HIV. Cell Host Microbe 23, 14–26 (2018).

8. Thorlund, K., Horwitz, M. S., Fife, B. T., Lester, R. & Cameron, D. W. Landscape review of current HIV ‘kick and kill’ cure research - some kicking, not enough killing. BMC Infect Dis 17, 595 (2017).

9. Julg, B. et al. Safety and antiviral effect of a triple combination of HIV-1 broadly neutralizing antibodies: a phase 1/2a trial. Nat Med 30, 3534–3543 (2024).

10. Bailón, L. et al. Safety, immunogenicity and effect on viral rebound of HTI vaccines combined with a TLR7 agonist in early-treated HIV-1 infection: a randomized, placebo-controlled phase 2a trial. Nat Commun 16, 2146 (2025).

11. Gurrola, T. E. et al. Delivering CRISPR to the HIV-1 reservoirs. Front Microbiol 15, 1393974 (2024).

12. Beliakova-Bethell, N. Eliminating the persistent HIV reservoir based on biomarker expression – How do we get there? Virology 603, 110368 (2025).

13. Magro, G., Calistri, A. & Parolin, C. How to break free: HIV-1 escapes from innovative therapeutic approaches. Front. Virol. 2, (2022).

14. Tee, K. K., Thomson, M. M. & Hemelaar, J. Editorial: HIV-1 genetic diversity, volume II. Front. Microbiol. 13, (2022).

15. Deng, K. et al. Broad CTL response is required to clear latent HIV-1 due to dominance of escape mutations. Nature 517, 381–385 (2015).

16. Cheeseman, H. M. et al. Broadly Neutralizing Antibodies Display Potential for Prevention of HIV-1 Infection of Mucosal Tissue Superior to That of Nonneutralizing Antibodies. Journal of Virology 91, (2017).

17. Salgado, M. et al. HLA-B*57 Elite Suppressor and Chronic Progressor HIV-1 Isolates Replicate Vigorously and Cause CD4+ T Cell Depletion in Humanized BLT Mice. J Virol 88, 3340–3352 (2014).

18. Hartana, C. A. et al. IL-15-dependent immune crosstalk between natural killer cells and dendritic cells in HIV-1 elite controllers. Cell Rep 42, 113530 (2023).

19. Fritschi, C. J. et al. Indoline CD4-mimetic compounds mediate potent and broad HIV-1 inhibition and sensitization to antibody-dependent cellular cytotoxicity. Proc Natl Acad Sci U S A 120, e2222073120.

20. van Aalst, S. et al. Routing dependent immune responses after experimental R848-adjuvated vaccination. Vaccine 36, 1405–1413 (2018).

21. Xiao, T., Cai, Y. & Chen, B. HIV-1 Entry and Membrane Fusion Inhibitors. Viruses 13, 735 (2021).

22. Itell, H. L., Humes, D. & Overbaugh, J. Several cell-intrinsic effectors drive type I interferon-mediated restriction of HIV-1 in primary CD4+ T cells. Cell Reports 42, (2023).

23. Stunnenberg, M., van Hamme, J. L., Zijlstra-Willems, E. M., Gringhuis, S. I. & Geijtenbeek, T. B. H. Crosstalk between R848 and abortive HIV-1 RNA-induced signaling enhances antiviral immunity. J Leukoc Biol 112, 289–298 (2022).

24. Utay, N. S. & Douek, D. C. Interferons and HIV Infection: The Good, the Bad, and the Ugly. Pathog Immun 1, 107–116 (2016).

25. van Broekhoven, C. L., Parish, C. R., Demangel, C., Britton, W. J. & Altin, J. G. Targeting dendritic cells with antigen-containing liposomes: a highly effective procedure for induction of antitumor immunity and for tumor immunotherapy. Cancer Res 64, 4357–4365 (2004).

26. Fox, C. B. et al. A nanoliposome delivery system to synergistically trigger TLR4 AND TLR7. Journal of Nanobiotechnology 12, 17 (2014).

27. Scheepers, M. R. W., IJzendoorn, L. J. van & Prins, M. W. J. Multivalent weak interactions enhance selectivity of interparticle binding. PNAS 117, 22690–22697 (2020).

28. Deak, P. E., Vrabel, M. R., Kiziltepe, T. & Bilgicer, B. Determination of Crucial Immunogenic Epitopes in Major Peanut Allergy Protein, Ara h2, via Novel Nanoallergen Platform. Scientific Reports 7, 3981 (2017).

29. Stefanick, J. F., Ashley, J. D., Kiziltepe, T. & Bilgicer, B. A Systematic Analysis of Peptide Linker Length and Liposomal Polyethylene Glycol Coating on Cellular Uptake of Peptide-Targeted Liposomes. ACS Nano 7, 2935–2947 (2013).

30. Jonak, Z. L. et al. A human lymphoid recombinant cell line with functional human immunodeficiency virus type 1 envelope. AIDS Res Hum Retroviruses 9, 23–32 (1993).

31. Vödrös, D. et al. Quantitative Evaluation of HIV-1 Coreceptor Use in the GHOST(3) Cell Assay. Virology 291, 1–11 (2001).

32. Navarro-Lopez, R. A., Silva-Campa, E., Santos-López, G. & Garibay-Escobar, A. Development and validation of a dynamic light scattering-based method for viral quantification: A straightforward protocol for a demanding task. PLOS ONE 20, e0324298 (2025).

33. Aneja, R. et al. Peptide Triazole Inactivators of HIV-1 Utilize a Conserved Two-Cavity Binding Site at the Junction of the Inner and Outer Domains of Env gp120. J. Med. Chem. 58, 3843–3858 (2015).

34. Chain, C. Y., Daza Millone, M. A., Cisneros, J. S., Ramirez, E. A. & Vela, M. E. Surface Plasmon Resonance as a Characterization Tool for Lipid Nanoparticles Used in Drug Delivery. Front. Chem. 8, (2021).

35. Wu, X. et al. Rational Design of Envelope Identifies Broadly Neutralizing Human Monoclonal Antibodies to HIV-1. Science 329, 856–861 (2010).

36. Browning, J. et al. Mice transgenic for human CD4 and CCR5 are susceptible to HIV infection. Proc Natl Acad Sci U S A 94, 14637–14641 (1997).

37. Banchereau, J. et al. Immunobiology of Dendritic Cells. Annual Review of Immunology 18, 767–811 (2000).

38. Ramakrishna, V. et al. Toll-like receptor activation enhances cell-mediated immunity induced by an antibody vaccine targeting human dendritic cells. Journal of Translational Medicine 5, 5 (2007).

39. Biancotto, A. et al. HIV-1–induced activation of CD4+ T cells creates new targets for HIV-1 infection in human lymphoid tissue ex vivo. Blood 111, 699–704 (2008).

40. McClelland, R. D., Culp, T. N. & Marchant, D. J. Imaging Flow Cytometry and Confocal Immunofluorescence Microscopy of Virus-Host Cell Interactions. Front Cell Infect Microbiol 11, 749039 (2021).

41. Costes, S. V. et al. Automatic and quantitative measurement of protein-protein colocalization in live cells. Biophys J 86, 3993–4003 (2004).

42. Soldemo, M., Pedersen, G. K. & Hedestam, G. B. K. HIV-1 Env-Specific Memory and Germinal Center B Cells in C57BL/6 Mice. Viruses 6, 3400–3414 (2014).

43. Mitchell, J. L. et al. Plasmacytoid dendritic cells sense HIV replication before detectable viremia following treatment interruption. J Clin Invest 130, 2845–2858 (2020).

44. Musumeci, A., Lutz, K., Winheim, E. & Krug, A. B. What Makes a pDC: Recent Advances in Understanding Plasmacytoid DC Development and Heterogeneity. Frontiers in Immunology 10, (2019).

45. Lichterfeld, M. et al. HIV-1–specific cytotoxicity is preferentially mediated by a subset of CD8+ T cells producing both interferon-γ and tumor necrosis factor–α. Blood 104, 487–494 (2004).

46. Lai, R. P. J. et al. Nef Decreases HIV-1 Sensitivity to Neutralizing Antibodies that Target the Membrane-proximal External Region of TMgp41. PLOS Pathogens 7, e1002442 (2011).

47. Collins, D. R., Gaiha, G. D. & Walker, B. D. CD8+ T cells in HIV control, cure and prevention. Nat Rev Immunol 20, 471–482 (2020).

48. Deng, Z., Yan, H., Lambotte, O., Moog, C. & Su, B. HIV controllers: hope for a functional cure. Front. Immunol. 16, (2025).

49. Yucha, R. et al. Higher HIV-1 Env gp120-Specific Antibody-Dependent Cellular Cytotoxicity (ADCC) Activity Is Associated with Lower Levels of Defective HIV-1 Provirus. Viruses 15, 2055 (2023).

50. Tettamanti Boshier, F. A., et al. Substantial uneven proliferation of CD4+ T cells during recovery from acute HIV infection is sufficient to explain the observed expanded clones in the HIV reservoir. Journal of Virus Eradication 8, 100091 (2022).

51. Stano, A. et al. Dense Array of Spikes on HIV-1 Virion Particles. J Virol 91, e00415–17 (2017).

52. Gray, E. R. et al. p24 revisited: A landscape review of antigen detection for early HIV diagnosis. AIDS 32, 2089–2102 (2018).

53. Tenchov, R., Bird, R., Curtze, A. E. & Zhou, Q. Lipid Nanoparticles─From Liposomes to mRNA Vaccine Delivery, a Landscape of Research Diversity and Advancement. ACS Nano 15, 16982–17015 (2021).

54. Adugna, A. Therapeutic strategies and promising vaccine for hepatitis C virus infection. Immun Inflamm Dis 11, e977 (2023).

55. Ding, S. et al. VE607 stabilizes SARS-CoV-2 Spike in the “RBD-up” conformation and inhibits viral entry. iScience 25, 104528 (2022).

56. Ali, M. M., Karasneh, G. A., Jarding, M. J., Tiwari, V. & Shukla, D. A 3-O-Sulfated Heparan Sulfate Binding Peptide Preferentially Targets Herpes Simplex Virus 2-Infected Cells. J Virol 86, 6434–6443 (2012).

57. Gillgrass, A., Wessels, J. M., Yang, J. X. & Kaushic, C. Advances in Humanized Mouse Models to Improve Understanding of HIV-1 Pathogenesis and Immune Responses. Frontiers in Immunology 11, (2021).

58. McCormack, M. J. et al. Retrovirus-based pseudotyped virus neutralisation assays overestimate neutralising activity in sera from participants receiving integrase inhibitors. Sci Rep 15, 28580 (2025).

59. Cheng, L. et al. Human innate responses and adjuvant activity of TLR ligands in vivo in mice reconstituted with a human immune system. Vaccine 35, 6143–6153 (2017).

60. Bolte, S. & Cordelières, F. P. A guided tour into subcellular colocalization analysis in light microscopy. J Microsc 224, 213–232 (2006).

