## supplemental material and methods for "HIV Virion Capturing Liposomes for Therapeutic Vaccination"

**Supplemental Materials and Methods for HIV Virion Capturing Liposomes for Therapeutic Vaccination**

#Author has passed away prior to publication

**Synthesis of DY-IV-040**

*General Procedures*

All solvents were reagent or high-performance liquid chromatography (HPLC) grade. Anhydrous $CH_2Cl_2$ and THF were obtained from the Pure SolveTM PS-400 system under argon atmosphere. All reagents were purchased from commercially available sources and sued as received. Reactions were magnetically stirred under a nitrogen or argon atmosphere, unless otherwise noted and reactions were monitored by Thin layer chromatography (TLC) was performed on pre-coated silica gel 60 F-254 plates (40-55 micron, 230-400 mesh) and visualized by UV light. Yields refer to chromatographically and spectroscopically pure compounds. Optical rotations were measured on a JASCO P-2000 polarimeter. Proton ($^1$H) and carbon ($^{13}$C) NMR spectra were recorded on a Bruker Avance III 500-MHz spectrometer or a Bruker NEO600 600-MHz spectrometer. Chemical shifts (δ) are reported in parts per million (ppm) relative to chloroform (δ 7.26), or methanol (δ 3.31) for $^1$H NMR, and chloroform (δ 77.2) or methanol (δ 49.15) for $^{13}$C NMR. High resolution mass spectra (HRMS) were recorded at the University of Pennsylvania Mass Spectroscopy Service Center on either a VG Micromass 70/70H or VG ZAB-E spectrometer. Analytical HPLC was performed with a Waters HPLC-MS system consisting of a 515 pump and Sunfire C18 reverse phase column (20 µL injection volume, 5 µm packing material, 4.5 x 50 mm column dimensions) with detection accomplished by a Micromass ZQ mass spectrometer and 2996 PDA detector. Preparative-scale HPLC was carried out on a Waters AutoPurification system (Milford, MA) equipped with a 3100 mass detector, a 2767 sample manager and a 2489 UV/visible detector. Purification was done on a 19x100mm SunFire® Prep C18 OBD 5µm column using a binary solvent gradient with mobile phase A (0.1% formic acid in water) and B (0.1% formic acid in acetonitrile). HPLC grade water and acetonitrile as well as Optima™ LC/MS-grade formic acid were purchased from Fisher

Scientific and used without further purification. Fraction collection was triggered using the mass detector. The purity of new compounds was judged by NMR and LCMS (>95%).

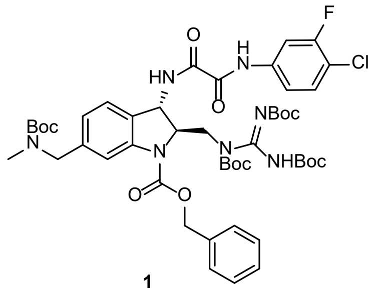

1

**benzyl (2R,3S)-6-(((tert-butoxycarbonyl)(methyl)amino)methyl)-3-(2-((4-chloro-3-fluorophenyl)amino)-2-oxoacetamido)-2-(((E)-1,2,3-tris(tert-butoxycarbonyl)guanidino)methyl)indoline-1-carboxylate (1):** Was synthesized according to literature procedure, and spectral data was compared to known references to confirm identity of the product.[1]

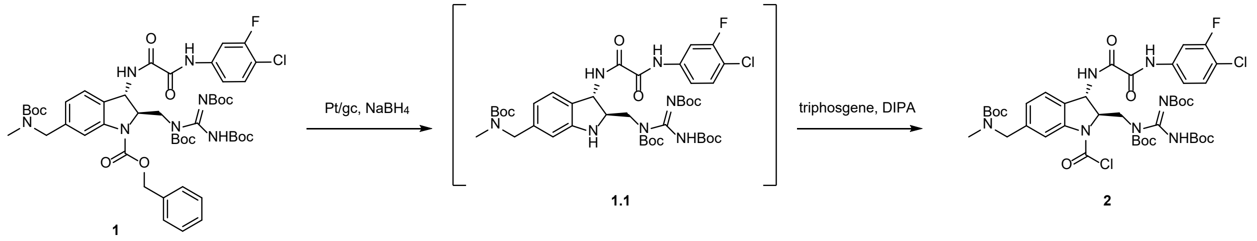

**(2R,3S)-6-(((tert-butoxycarbonyl)(methyl)amino)methyl)-3-(2-((4-chloro-3-fluorophenyl)amino)-2-oxoacetamido)-2-(((E)-1,2,3-tris(tert-butoxycarbonyl)guanidino)methyl)indoline-1-carbonyl chloride (2):** Compound (**1**) (200 mg, 0.204 mmol, 1.0 equiv) was added to a microwave vial followed by platinum on graphitized carbon, 20% loading (20 mg, 10% w/w), and anhydrous methanol (2.04 mL, 10ml/mmol). Subsequently, $NaBH_4$ (8.48 mg, 0.224 mmol, 1.1 equiv) was added and an aluminum crimp seal top was added on top of the microwave vial and immediately sealed. Upon addition of the $NaBH_4$ the solution bubbled indicated production of $H_2$ (g). This vessel was transferred to an aluminum block and heated at 40 °C for 1 hour, at which time UPLCMS indicated complete consumption of the starting material. The reaction was then diluted in methanol and filtered through a pad of celite, washed with methanol (2 mL) and concentrated *in vacuo*. The concentrated product was once again taken up in EtOAc (2 mL), and extracted with water (2 x 2 mL), washed with brine (2 mL), dried over $Na_2SO_4$, and concentrated *in vacuo* to yield a white solid. The crude material was then immediately passed through a plug of silica pre-treated with 10% EtOAc/hexanes with a 1%

[1] Fritschi, C. J., Anang, S., Gong, Z., Mohammadi, M., Richard, J., Bourassa, C., ... & Smith III, A. B. (**2023**). Indoline CD4-mimetic compounds mediate potent and broad HIV-1 inhibition and sensitization to antibody-dependent cellular cytotoxicity. *Proceedings of the National Academy of Sciences*, *120*(13), e2222073120.

$NEt_3$ buffer and eluted with the same solvent system until elution of the desired product **1.1** was observed by TLC (142 mg, 84% yield).

The material (**1.1**) was then redissolved in anhydrous $CH_2Cl_2$ (0.84 mL, 5 mL/mmol), followed by the addition of diisopropylamine (0.0354 mL, 0.251 mmol, 1.5 equiv). Separately, triphosgene (17.35 mg, 0.058 mmol, 0.15 equiv) was dissolved in anhydrous $CH_2Cl_2$ (0.290 mL, 5 mL/mmol). The triphosgene solution was then added to the starting material solution dropwise at 0 °C and stirred for 3 hours at which time UPLCMS analysis indicated consumption of the starting material. The reaction was then quenched with $NH_4Cl$ (1 mL), and the aqueous layer was extracted with $CH_2Cl_2$ (3 x 2 mL). The organic layers were separated, dried with $Na_2SO_4$, filtered and concentrated *in vacuo*. The crude carbamoyl chloride (**2**) was then purified by flash chromatography (20% EtOAc/hexanes) to yield **2** as a white powder (160 mg, 83% yield over 2 steps).

**$^1$H NMR (500 MHz, MeOD)** δ 7.83 – 7.80 (bs, 1H), 7.82 (dd, *J* = 11.4, 2.4 Hz, 1H), 7.53 – 7.29 (m, 4H), 5.48 – 5.18 (m, 2H), 4.44 (d, *J* = 17.7 Hz, 2H), 4.18 – 4.06 (m, 1H), 3.97 (dt, *J* = 14.7, 8.2 Hz, 1H), 2.83 (s, 3H), 1.56 – 1.39 (m, 36H).

**$^{13}$C NMR (126 MHz, MeOD)** δ 160.92, 160.08 (d, *J* = 245.22 Hz), 159.62, 154.86, 146.32, 139.04 (d, *J* = 9.89 Hz), 131.66, 129.61, 118.05, 117.05 (d, *J* = 17.41 Hz), 109.60 (d, *J* = 25.89 Hz), 85.92, 61.54, 53.66, 53.16, 28.73, 28.47, 28.30, 20.86, 14.46.

**HRMS** (ESI) *m/z* [M+H]+ calcd for $C_{41}H_{55}Cl_2FN_7O_{11}$ : 910.3321, found 910.3319;

**$[\alpha]_D^{24}$** +68.2 (*c* 0.50, MeOH).

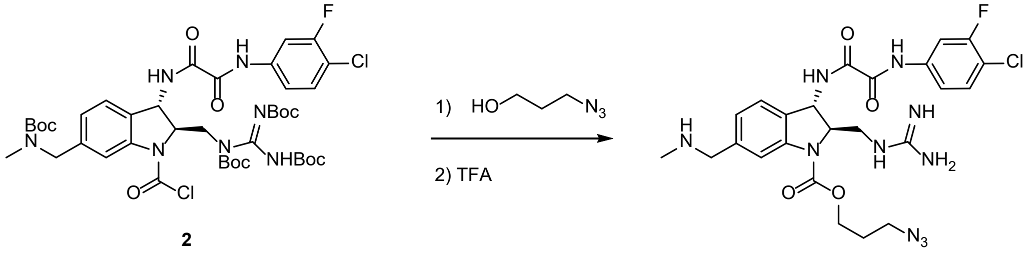


**2**

**3**
**(DY-IV-040)**

**3-azidopropyl (2R,3S)-3-(2-((4-chloro-3-fluorophenyl)amino)-2-oxoacetamido)-2-(guanidinomethyl)-6-((methylamino)methyl)indoline-1-carboxylate (DY-IV-040) (3):** Compound **2** was dissolved in $CH_2Cl_2$ (0.1 M), followed by addition of 3-azidopropan-1-ol (1.1 equiv.). N,N-dimethylaminopyridine (1.2 equiv.) was then added in one portion, and the mixture was allowed to stir overnight, at which time UPLCMS analysis indicated consumption of **2**. The reaction was quenched with sat. aq. $NH_4Cl$, and the aqueous layer was extracted 3 x $CH_2Cl_2$. The organic layers were separated, dried with $Na_2SO_4$, filtered and concentrated *in vacuo*. The crude **3** (1.0 equiv.) was then taken up in $CH_2Cl_2$ (0.1 M) and cooled to 0 °C. Trifluoroacetic acid (28 equiv.) was added and the mixture was allowed to warm to rt. The reaction was allowed to stir for 18 hours at which time UPLCMS analysis indicated consumption of the starting material. Upon completion, solvent was removed *in vacuo* and the product was taken up in 1:1 MeCN/$H_2O$ and purified via Waters AutoPur with mass directed HPLC with a flow rate of 32 mL/min and the gradient program as follows: 0-0.5min 15% B, 1-9min linear from 15% to 35% B, 9-9.5min linear from 35% to 95% B, 9.5-11.5min 95% B and 11.5-12min 10% B. Fractions were collected based on a mass trigger and was lyophilized to yield the final product, **3 (DY-IV-040)** as a colorless oil.

**$^{1}$H NMR (500 MHz, MeOD)** δ 7.94 (bs, 1H), 7.83 (dd, *J* = 11.4, 2.2 Hz, 1H), 7.54 – 7.41 (m, 3H), 7.23 (dd, *J* = 7.8, 1.6 Hz, 1H), 5.24 (d, *J* = 2.4 Hz, 1H), 4.58 (bm, 1H), 4.45 (m, 1H), 4.23 (s, 2H), 3.63 – 3.47 (m, 4H), 2.73 (s, 3H), 2.04 (m, 2H);

**$^{13}$C NMR (126 MHz, MeOD)** δ 162.90 (q, $J_{CF}$ = 35.57 Hz, TFA), 161.31, 161.28, 159.94 (d, $J_{CF}$ = 246.54 Hz), 159.29, 159.23, 138.96 (d, $J_{CF}$ = 11.27 Hz), 134.57, 131.75, 128.00. 126.45, 118.31, 118.08 (d, $J_{CF}$ = 3.49 Hz), 117.41 (d, $J_{CF}$ = 18.13 Hz), 109.85 (d, $J_{CF}$ = 25.71 Hz), 66.71, 65.62, 54.44, 53.55, 49.57, 44.48, 33.11, 29.38;

**HRMS** (ESI) m/z 575.2043 [calcd for $C_{24}H_{29}ClFN_{10}O_4$ (M+H)+ 575.203

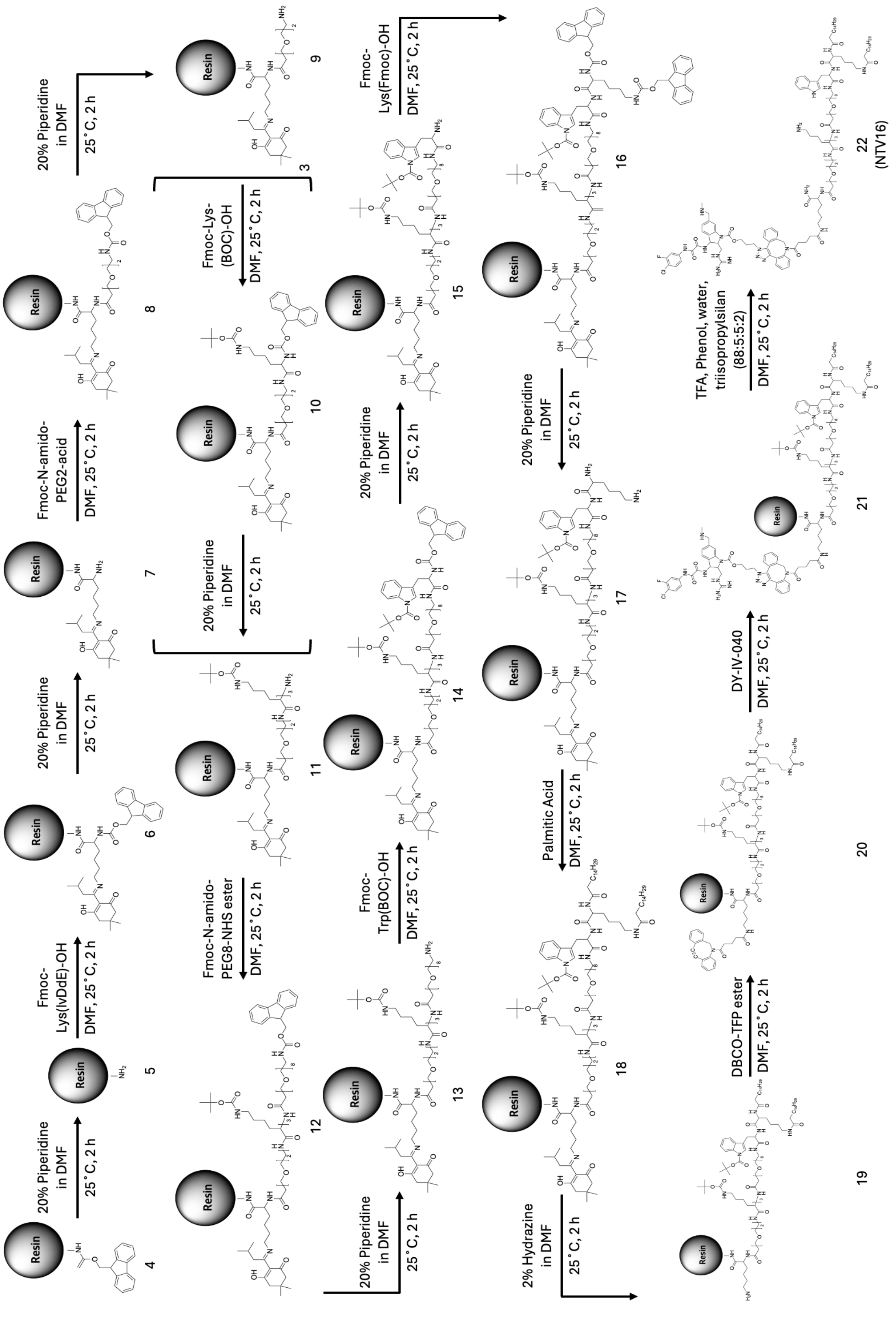


### CJF-lipids for Synthesis

*General Procedures*

Rink Amide MBHA resin (100–200 mesh), Fmoc-Lys(IvDde)-OH, Fmoc-Arg(Pbf)-OH, Fmoc-Trp(Boc)-OH, Fmoc-Lys(Fmoc)-OH, and piperidine were obtained from Sigma-Aldrich. HBTU, DIPEA, and anhydrous DMF (HPLC grade, 99.7%) were purchased from Thermo Scientific. Fmoc-N-amido-PEG2-acid was purchased from Quanta Biodesign, and Fmoc-Lys(Boc)-OH from Anaspec.
Fmoc-N-amido-PEG8-NHS ester and DBCO-TFP ester were obtained from Vector Labs.
Palmitic acid was purchased from Chem-Impex.
DCM, methanol, and TFA were purchased from VWR.
Hydrazine monohydrate (98%) and triisopropylsilane (TIS, 98%) were obtained from BeanTown Chemical.
HPLC-grade acetonitrile was purchased from Oakwood Chemical.

*Solid-Phase Synthesis of CJF-Lipid Precursors*

Solid-phase synthesis was carried out in a 25-mL Chemglass peptide synthesis vessel. Kaiser tests were used to confirm the presence of free amines after deprotection. All washing steps were performed with 3 mL of solvent unless otherwise note

Rink Amide MBHA resin **(4)** (0.01 mmol) was swollen in 10% DIPEA in DCM (3 mL) for 15 min on a flask shaker and washed with DCM. Fmoc deprotection was performed by adding 20% piperidine in DMF, shaking for 5 min, followed by three DCM washes. This deprotection/washing step was repeated twice to fully remove the Fmoc group, yielding resin **(5)**. Successful deprotection was confirmed using a Kaiser test. The resin was washed with DMF, DCM, and DMF (3× each), then 4 equiv Fmoc-N-amido-PEG2-acid (0.04 mmol) and 4 equiv HBTU (0.04 mmol) were dissolved in DMF (3 mL) and pre-cooled at –20 °C for 5 min. DIPEA (41.84 µL, 0.04 mmol) was added and the mixture was stirred for 5 min at room temperature before being transferred to the resin. The reaction was shaken for 2 h to afford **(6)**. For subsequent couplings up to intermediate **(18)**, the following cycle was repeated: washing → Kaiser test → Fmoc deprotection → washing → coupling of the next Fmoc-protected amino acid or PEG-derived building block. For coupling steps involving ester-functionalized compounds, HBTU was omitted. To remove the IvDde protecting group and generate **(19)**, the resin was treated with 2% hydrazine in DMF (3 mL) and shaken for 5 min at room temperature, followed by DCM washes and Kaiser testing. After confirming formation of **(19)**, the resin was washed (DMF/DCM/DMF) and treated with 4 equiv DBCO-TFP ester and 4 equiv DIPEA (0.04 mmol each) to yield **(20)** after 2 h shaking. Copper-free click chemistry was then performed by adding DY-IV-040 (3) (0.01 mmol) in DMF and shaking for 2 h at room temperature. The resin was washed with DMF (3×), then dried under nitrogen for 5 min to afford **(21)**.

*Solid-Phase Synthesis of CJF-Lipid Precursors*

Cleavage cocktail (Reagent B) was prepared as: TFA/phenol/water/TIS = 88:5:5:2 (v/v). Resin-bound **(21)** was treated with Reagent B (3 mL) at room temperature for 15 min to remove side-chain protecting groups and cleave the product from the resin. The crude NTV16-lipid **(22)** was collected into a round-bottom flask, and residual TFA was removed by rotary evaporation (IKA Works) at 50 °C and 50 mbar. Additional TFA was removed by a vacuum chamber. DI water was added to suspend the crude **(22)**, followed by freezing at −80 °C and lyophilization. The resulting powder was dissolved in a 1:1 mixture of DI water and acetonitrile and purified by preparative HPLC (Agilent) using a Polaris 5 C8 250 × 10 mm column. Purified **(22)** was characterized by MALDI-TOF (Bruker).

### NTV16 Formation

*General Procedures*

Chloroform (molecular biology grade) was purchased from MP Biomedicals.
DSPC (16:0), cholesterol (ovine wool, 98%), and 18:0 PC were obtained from Avanti.
DiD' oil was purchased from Invitrogen.
R848 (Resiquimod) was purchased from Chem-Impex.

*Preparation of NTV16 Liposomes*

Purified lipid **(22)** was dissolved in methanol/acetonitrile (1:1, v/v). For liposome formulation, DSPC (88%), cholesterol (5%), mPEG2000-DSPE (5%), DiD (1%), and **(22)** (1%) were dissolved in 200 µL chloroform. A thin lipid film was generated using a rotary evaporator. The dry film was hydrated with 200 µL of 166 µM R848 in PBS at 66 °C to allow R848 encapsulation. The resulting suspension was passed through a 100-nm polycarbonate membrane (Cytiva) using a mini-extruder. Seven extrusion cycles were performed to obtain uniformly sized liposomes. Unencapsulated R848 and unincorporated CJF-lipid were removed using Slide-A-Lyzer™ MINI dialysis devices (10 kDa MWCO). Final NTV16 particle size was measured by DLS (Brookhaven Instruments).
