## Supplemental Figure S1-16 for "HIV Virion Capturing Liposomes for Therapeutic Vaccination"

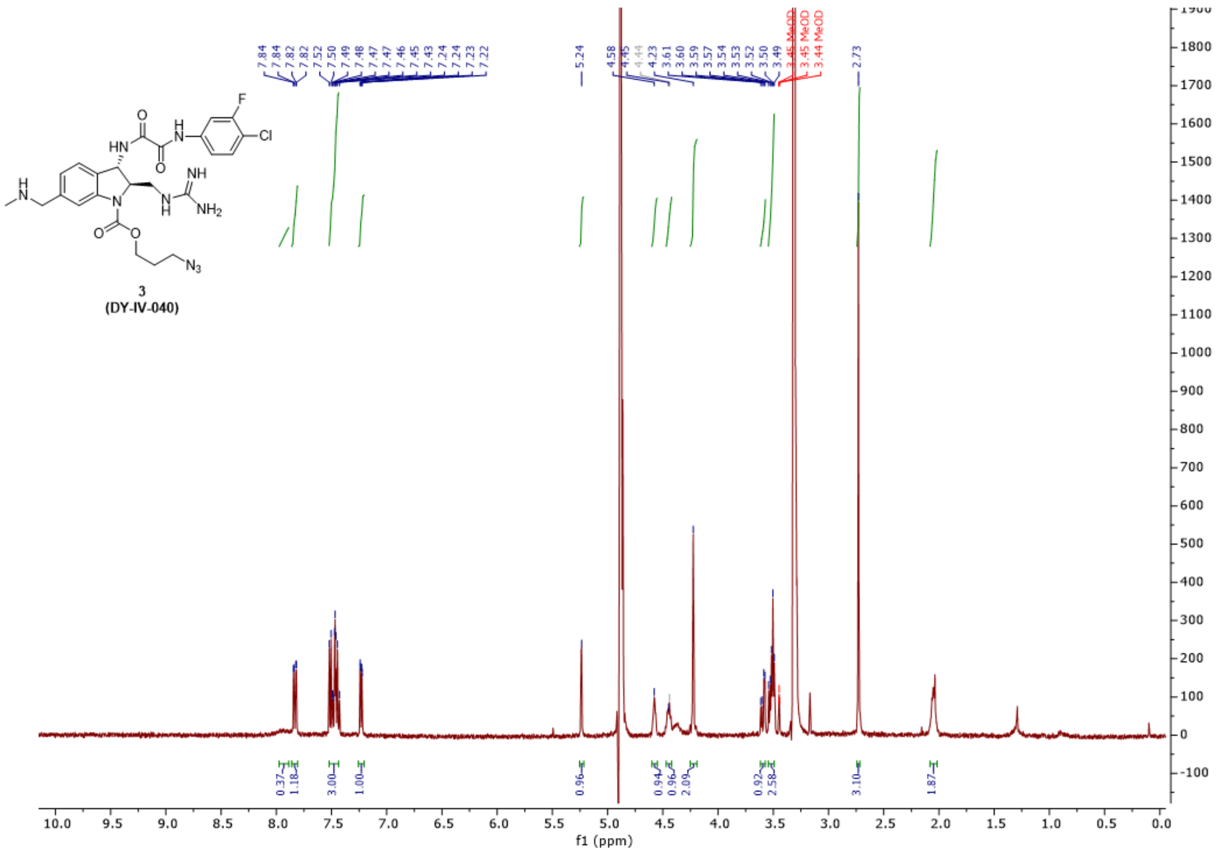


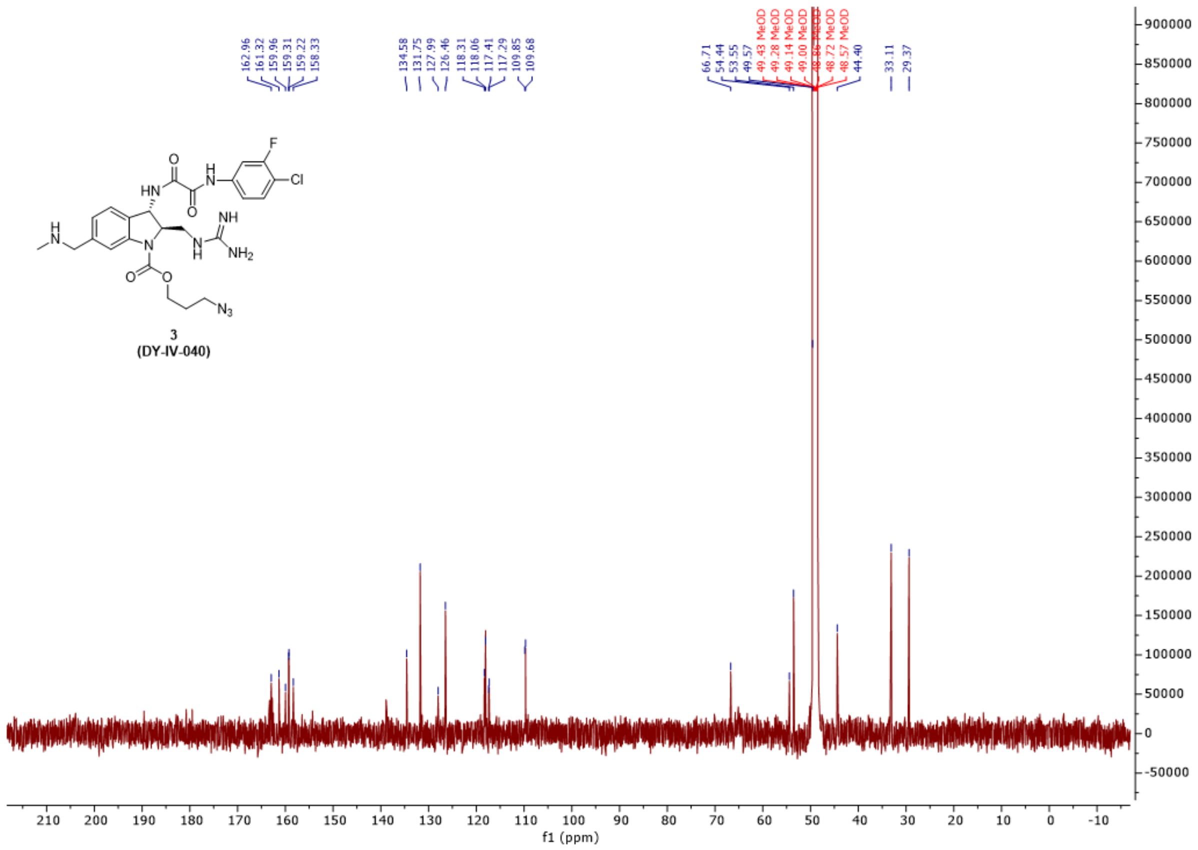


Figure S-1. H-NMR (top) and C-NMR (bottom) analysis of azide modified CJF-III-288 (DY-IV-040)

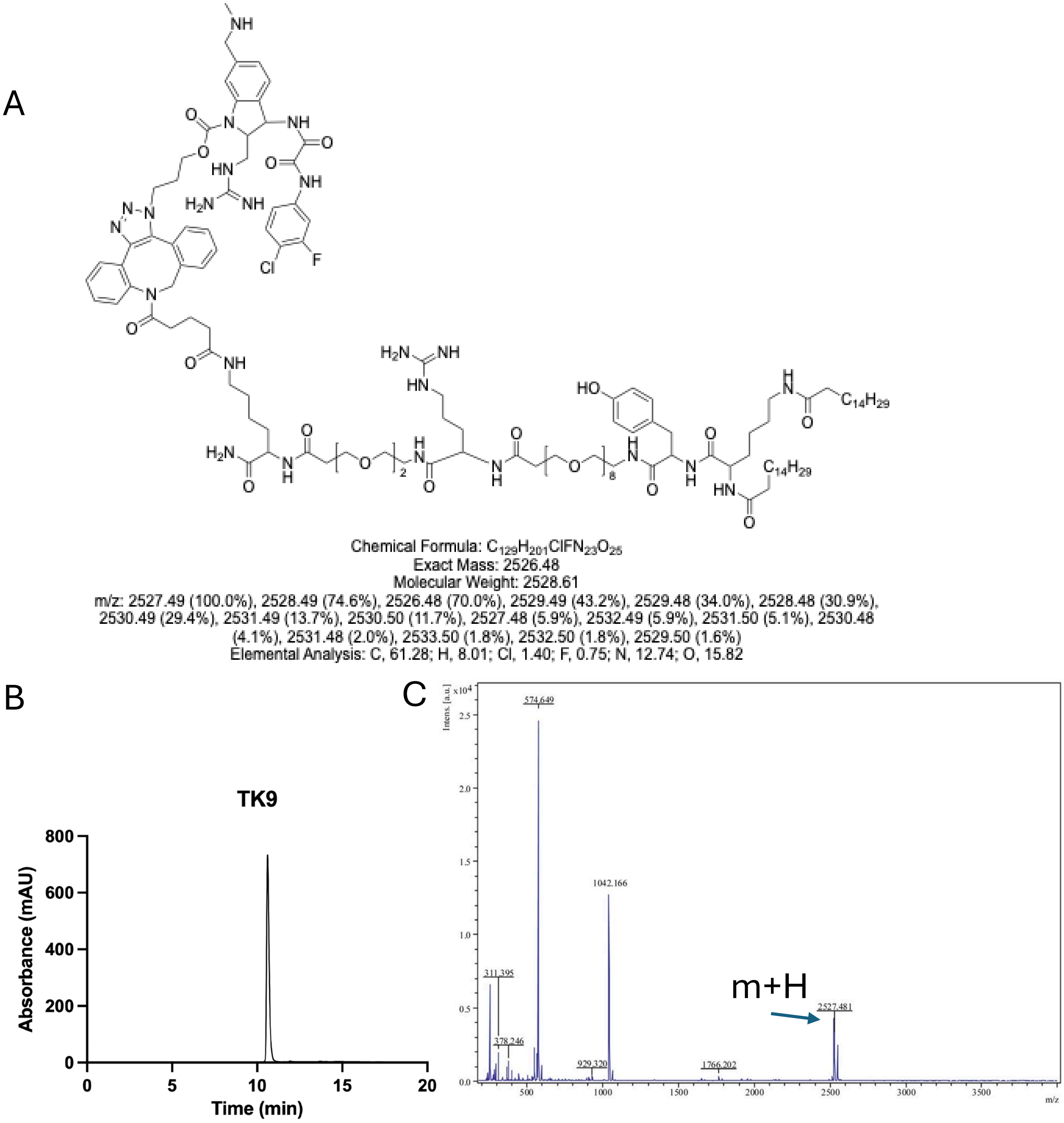


Figure S-2. Chemical Characterization of TK9 showing (A) chemical structure (B) HPLC analysis at 220 nm and (C) MALDI MS analysis confirming m+H peak.

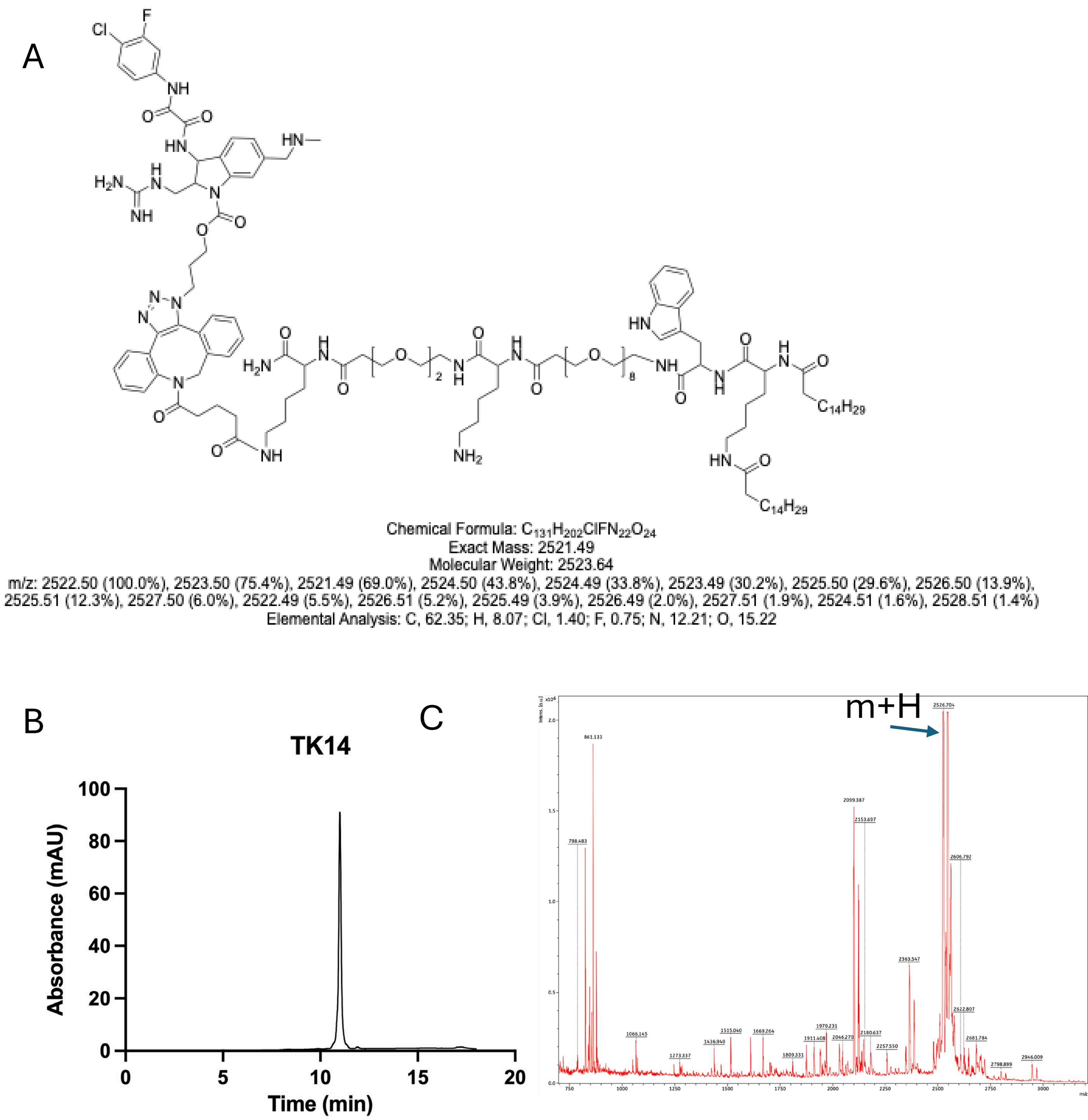


Figure S-3. Chemical Characterization of TK14 showing (A) chemical structure (B) HPLC analysis at 220 nm and (C) MALDI MS analysis confirming m+H peak.

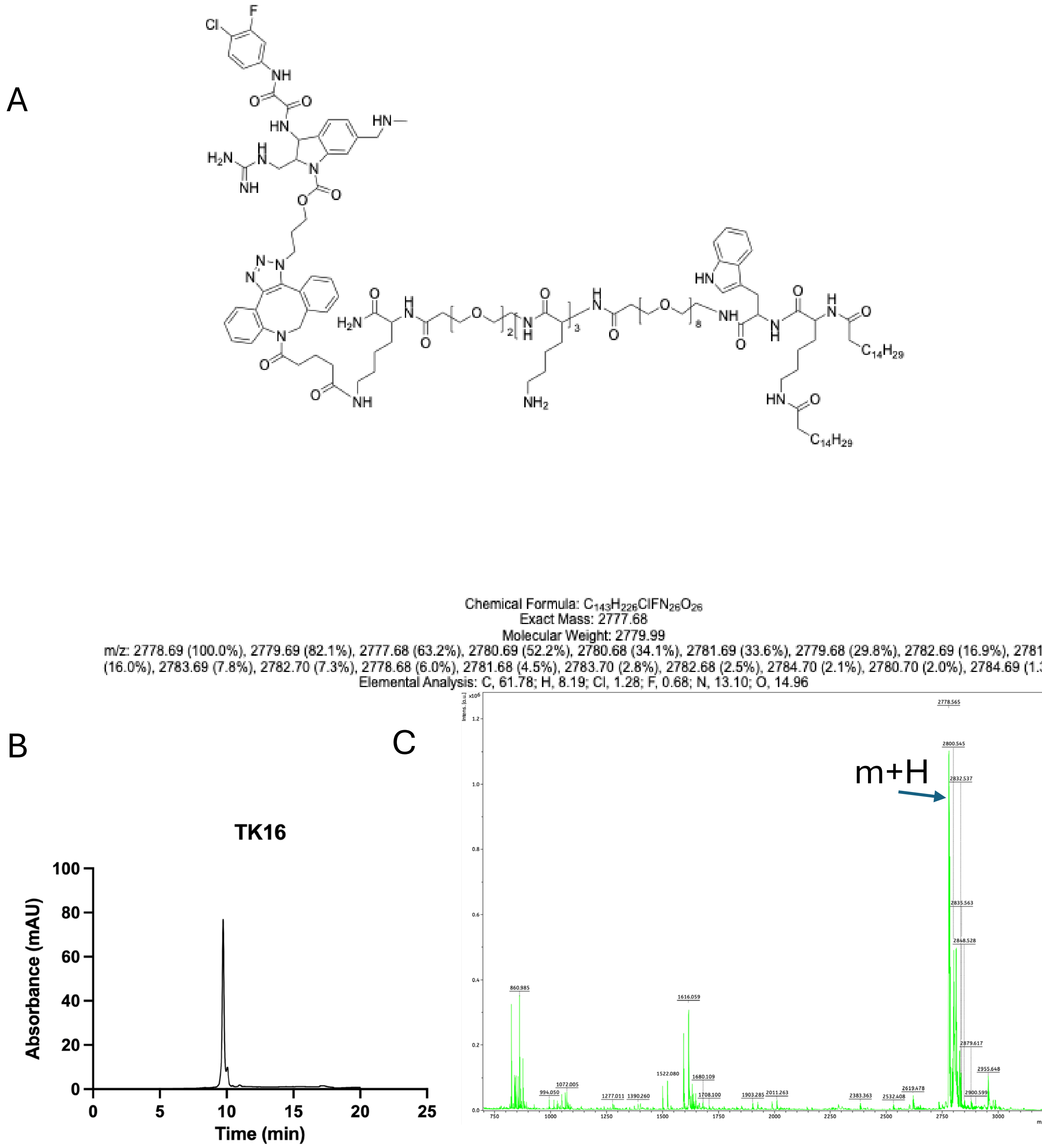


Figure S-4. Chemical Characterization of TK16 showing (A) chemical structure (B) HPLC analysis at 220 nm and (C) MALDI MS analysis confirming m+H peak.

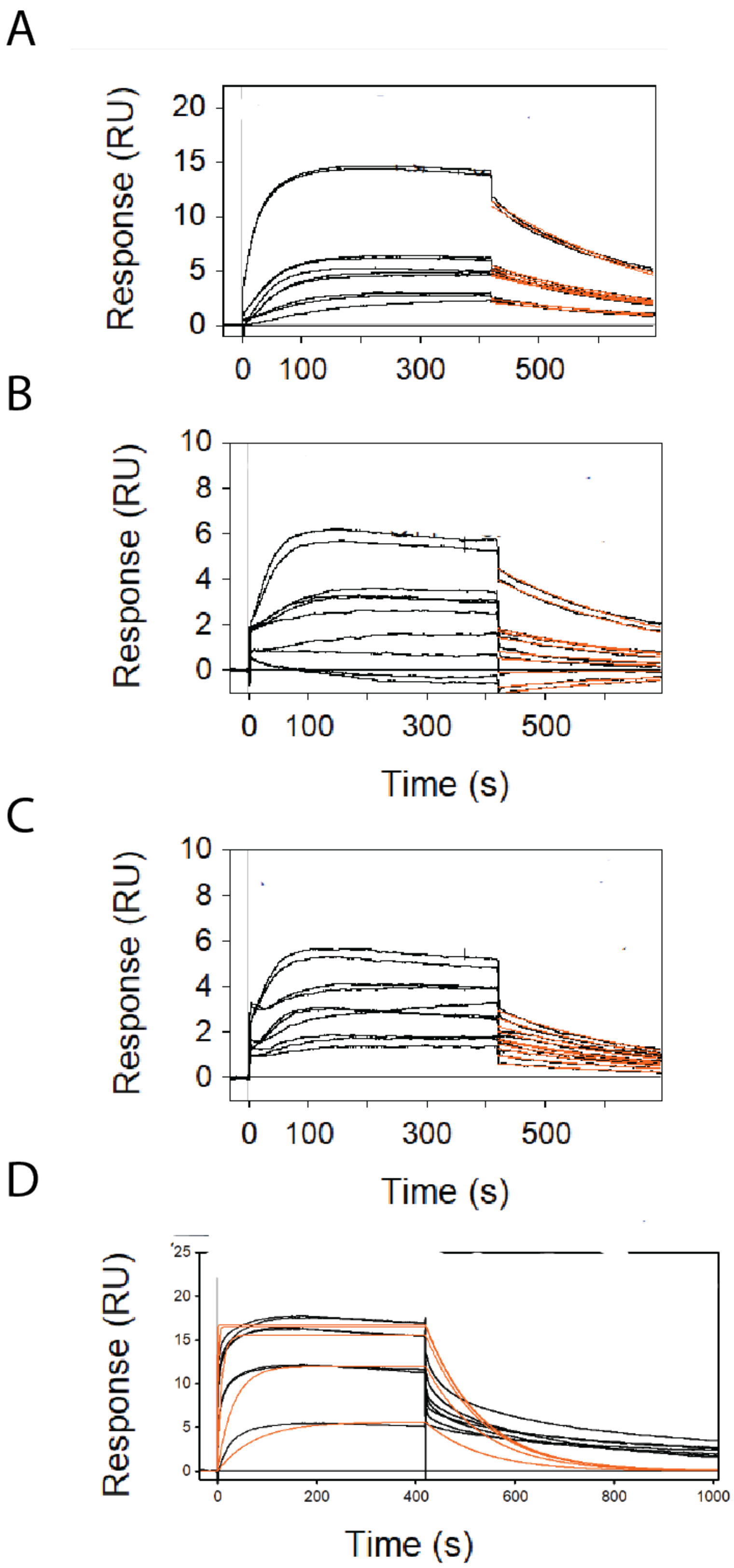


Figure S-5. Raw sensograms of data used in Figure 1E. SPR chips were coated with 8000 RFU of YU-2 gp120 protein then analyzed binding of 2000, 400, 80, 16 or 3.2 nM of various ligands (A) TK9, (B) TK14, (C)TK16 or (D) unmodified CJF-III-288. All samples were diluted in PBS-P with 2% DMSO and binding compared to buffer only control.

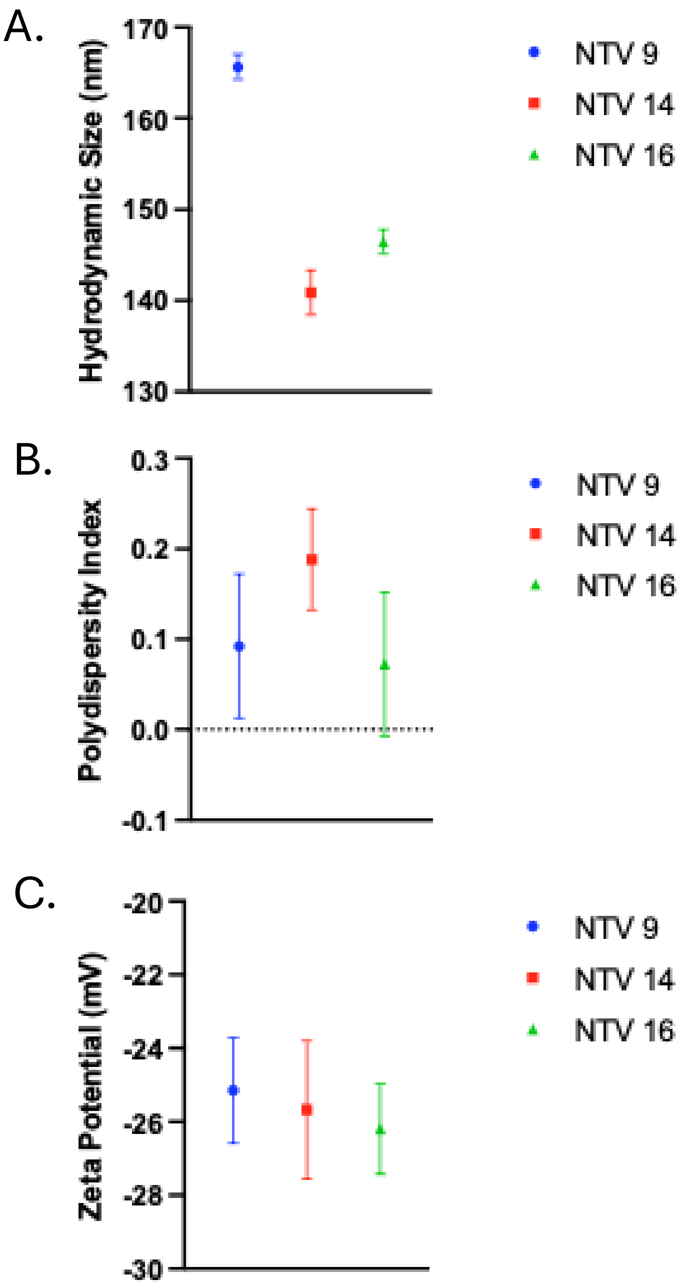


| Lipid Conjugate | Linker Type | Diameter (nM) | PDI | Zeta Potential (mV) |
|---|---|---|---|---|
| TK-9 | Arg | 166±1.4 | 0.09±0.08 | -25.1±1.4 |
| TK-14 | Lys | 141±2.4 | 0.19±0.05 | -25.7±1.9 |
| TK-16 | Lysx3 | 146±1.3 | 0.07±0.08 | -26.2±1.2 |

Figure S-6. DLS and zeta potential characterization of NTV formulation. Dynamic light scattering (DLS) was used to measure the (A) hydrodynamic diameter and (B) polydispersity index (PDI) of NTV-9, NTV-14, and NTV-16 formulations, while (C) electrophoretic light scattering was used to determine zeta potential. Formulations displayed comparable negative surface charge and low PDI values, indicating uniform particle distributions. Table summarizes mean diameter, PDI, and zeta potential for NTVs prepared with TK-9, TK-14, or TK-16 lipid conjugates.

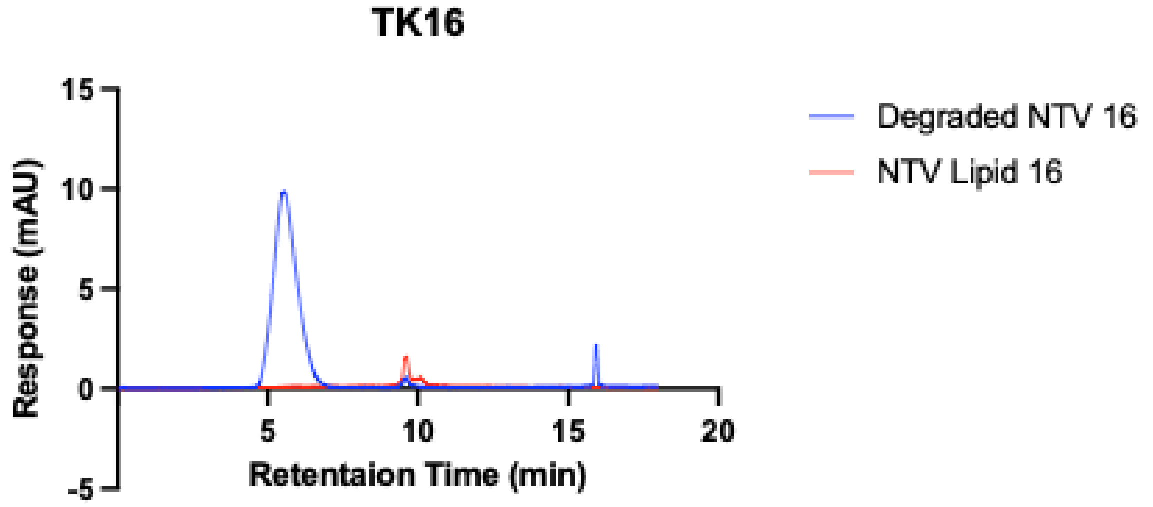


| Degraded NTV 16 (mAU) | NTV 16 Lipid (mAU) | Incorporation efficiency (%) |
|---|---|---|
| 0.5575 | 1.6187 | 34.44 |

Figure S-7. CJF-lipid encapsulation efficiency of NTV-16. Reversed-phase HPLC was used to quantify incorporation of NTV16 lipid into NTV formulations. To generate a fully released lipid reference, NTV-16 particles were degraded in acetonitrile (ACN) prior to injection (blue trace). Chromatograms of intact NTV-16 lipid (red trace) and ACN-degraded NTV-16 show the TK-16 lipid peak eluting at ~10 min. Peak heights (mAU) obtained from these lipid-associated signals were used to calculate incorporation efficiency, summarized in the table below. NTV-16 displayed an incorporation efficiency of 34.44%.

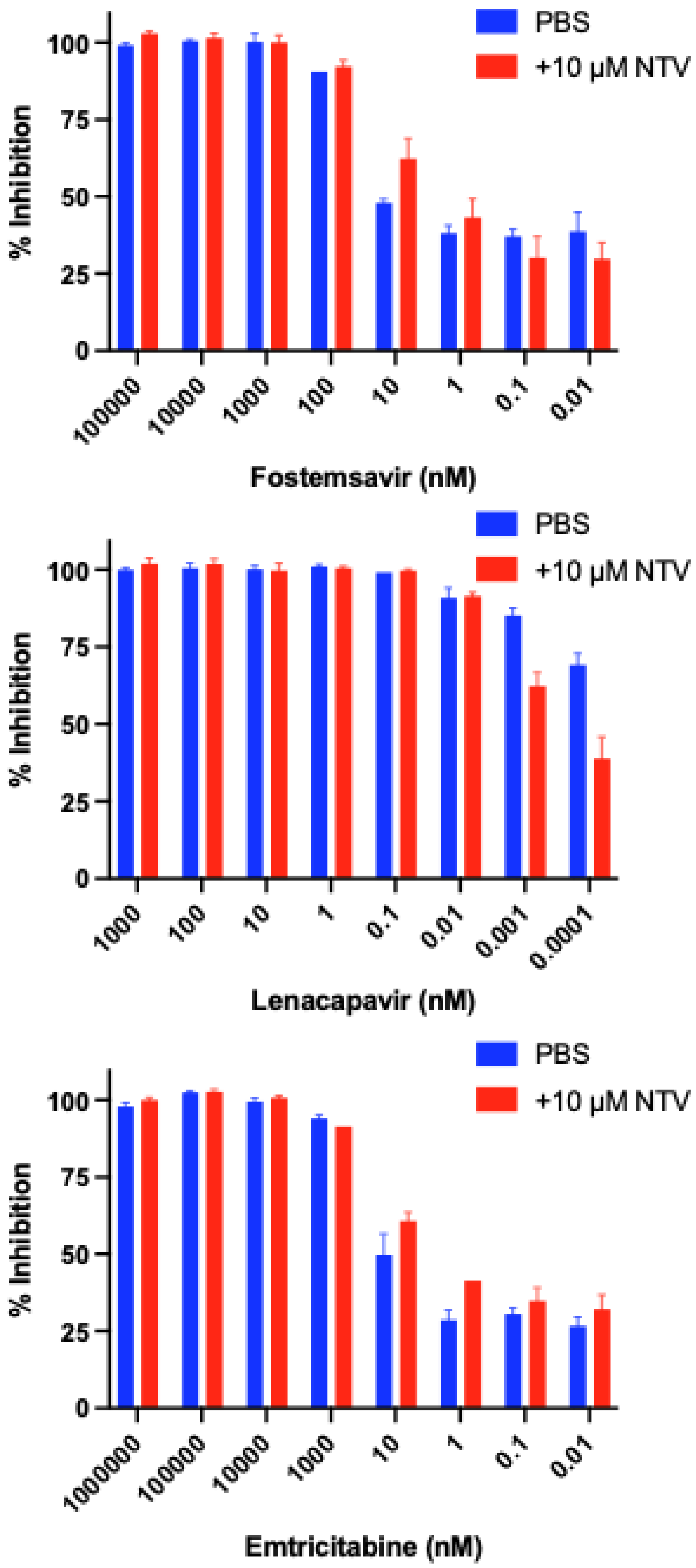


Figure S-8. Compatibility of NTV with antiretroviral drugs. HIV inhibition assays were performed to evaluate whether co-treatment with 10 µM NTV affects the antiviral activity of Fostemsavir, Lenacapavir, or Emtricitabine. Dose–response curves for each drug were measured in the presence of NTV (red) or PBS control (blue). Across all concentrations tested, NTV did not alter inhibition profiles or reduce drug potency, indicating no interference with ART activity. These results demonstrate that the NTV formulation is compatible with standard antiretroviral therapies.

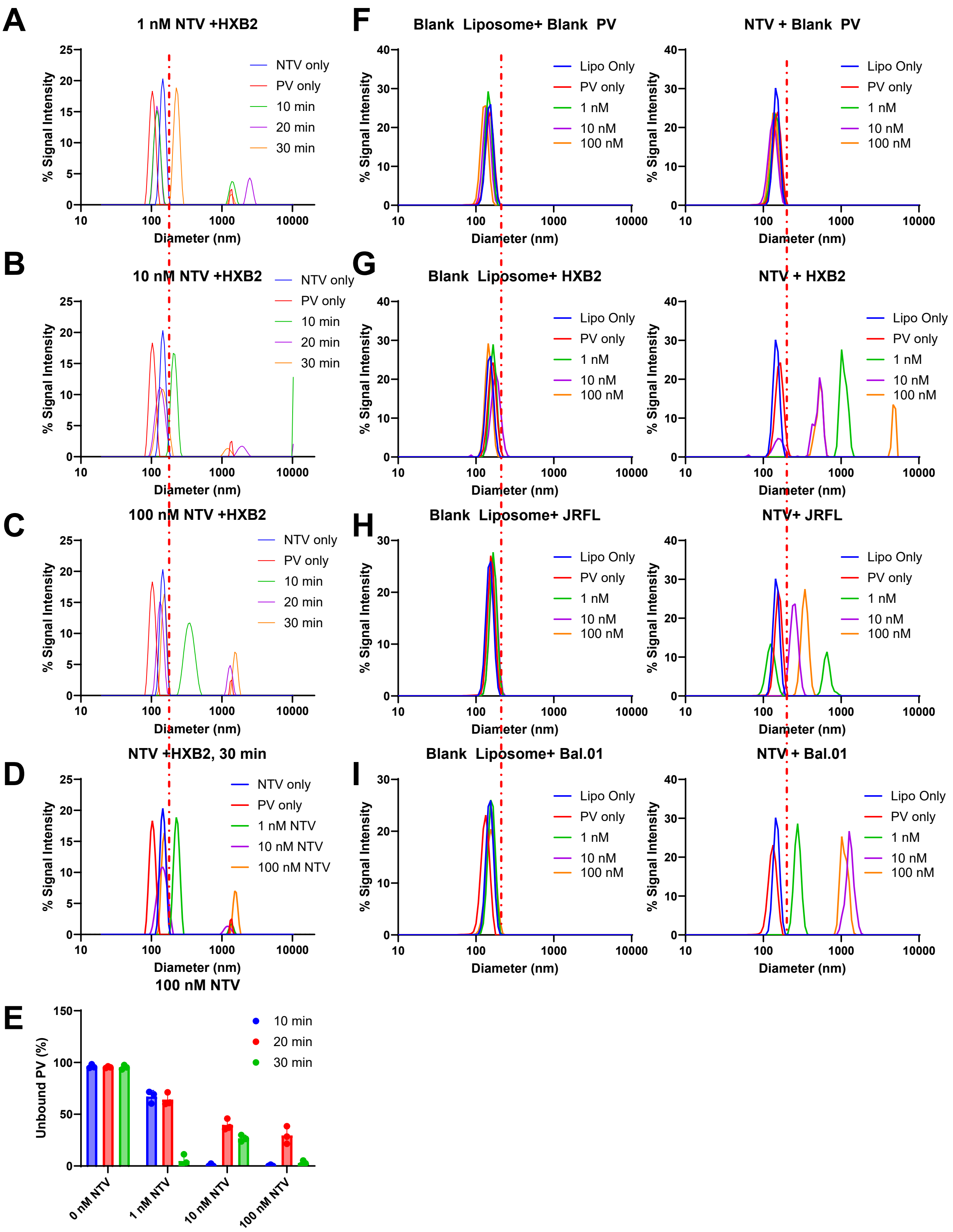


Figure S-9. Extended DLS Data for NTV-PV complex formation. (A-E) 0.5 ng/mL of DiO labeled HXB2 env PV was incubated with either 1% TK16 NTV 100 nm NTV or blank liposome and analyzed at 10, 20 or 30 minutes by SPR. Representative histogram plots for (A)1 nM total lipid NTV (B) 10 nM or (C) 100 nM. (D) Plots for 30 minutes for all concentration points. (E) Fraction unbound PV for triplicate measurements. (F-I) Similar to A-E, NTV (right) or blank liposome (left) were incubated for 30 minutes with varying env expressing PV, (F) env deficient PV, (G) HXB2, (H) JRFL or (I) Bal.01.Red line indicate cutoff for bound vs unbound PV. Error bars ±SD. PV only= no NTV control, NTV only= no PV control.

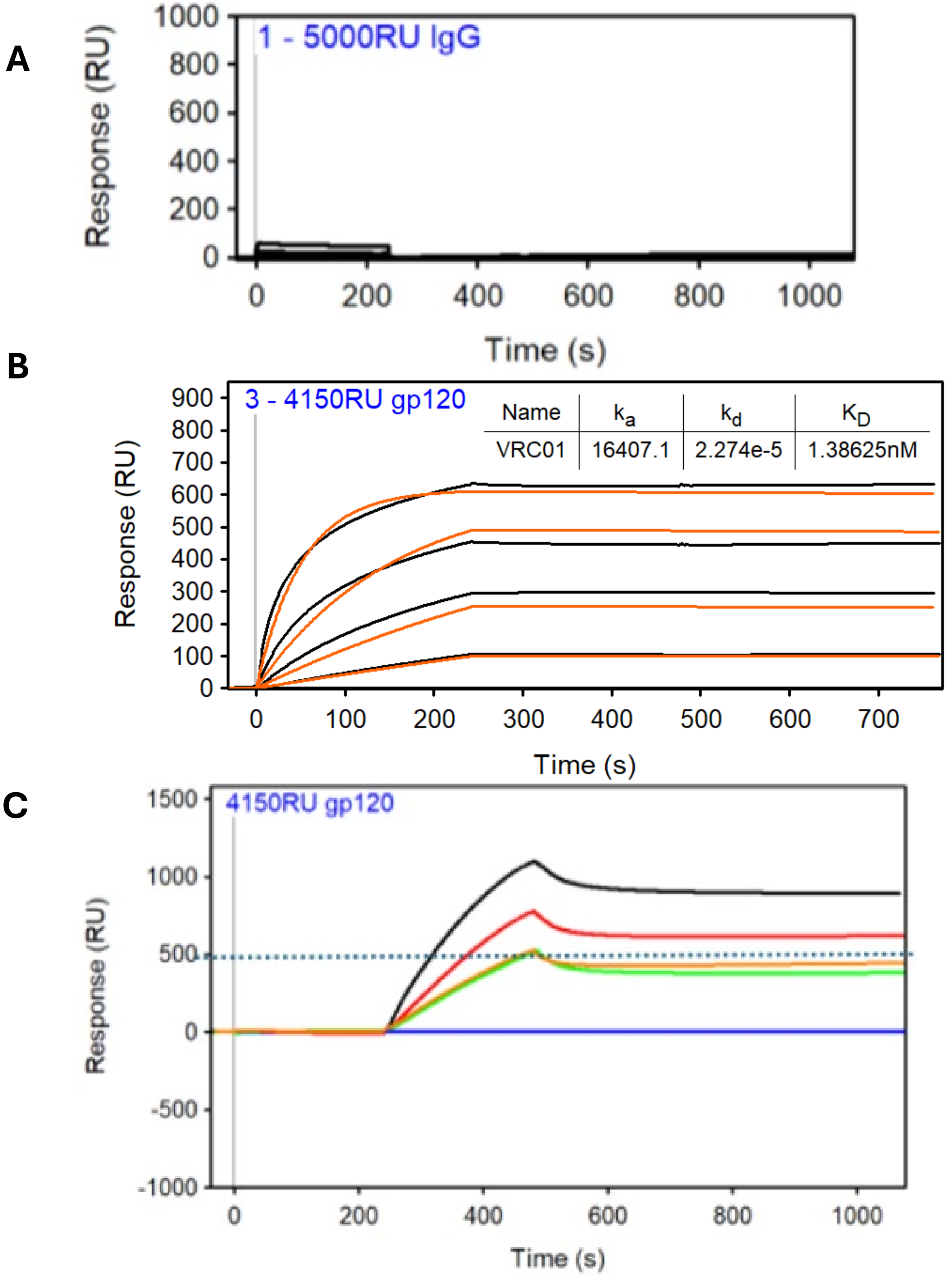


Figure S-10. Additional SPR data. (A) SPR of 100, 25, 6.25 μM total lipid of NTV (1% TK16, 100 nm) with SPR chip coated with 5000 RU of irrelevant control antibody. (B) Sensogram of 4150 RU of YU gp120 and 1200, 444, 133, or 44 nM of VCR01. (C) SPR chips treated in part B were then treated with 25 μM of NTV and response measured. Color indicate VCR01 antibody pretreated. Blank: 44 nM, Red: 133 nM, Orange: 444 nM and Green: 1200 nM. Fractional inhibition was calculated in Figure 2K.

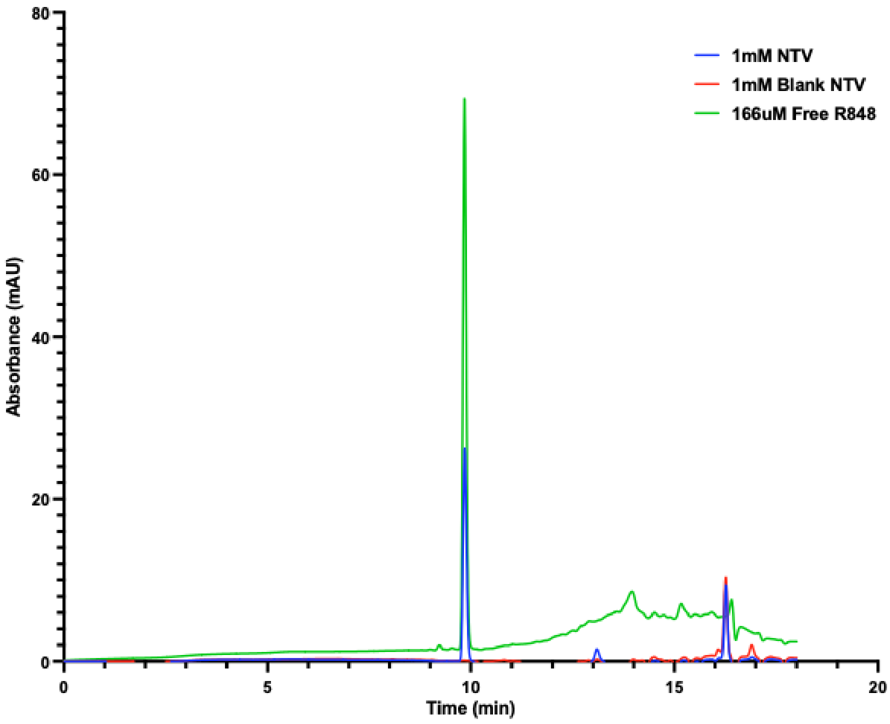


| 166uM R848 loaded1mM NTV (mAU) | 166uM Free R848 (mAU) | Incorporation efficiency (%) |
|---|---|---|
| 26.35 | 69.20 | 38.08 |

Figure S-11. R848 loading analysis of NTV by HPLC. Reversed-phase HPLC was used to quantify incorporation of R848 into 1 mM NTV formulations. Chromatograms of R848-loaded NTV (blue), blank NTV (red), and 166 μM free R848 (green) are shown, with R848 eluting at approximately 10 min. Peak intensities (mAU) at the R848 retention time were used to calculate loading efficiency. Comparison of the loaded NTV peak with the free R848 reference indicated an incorporation efficiency of 38%.

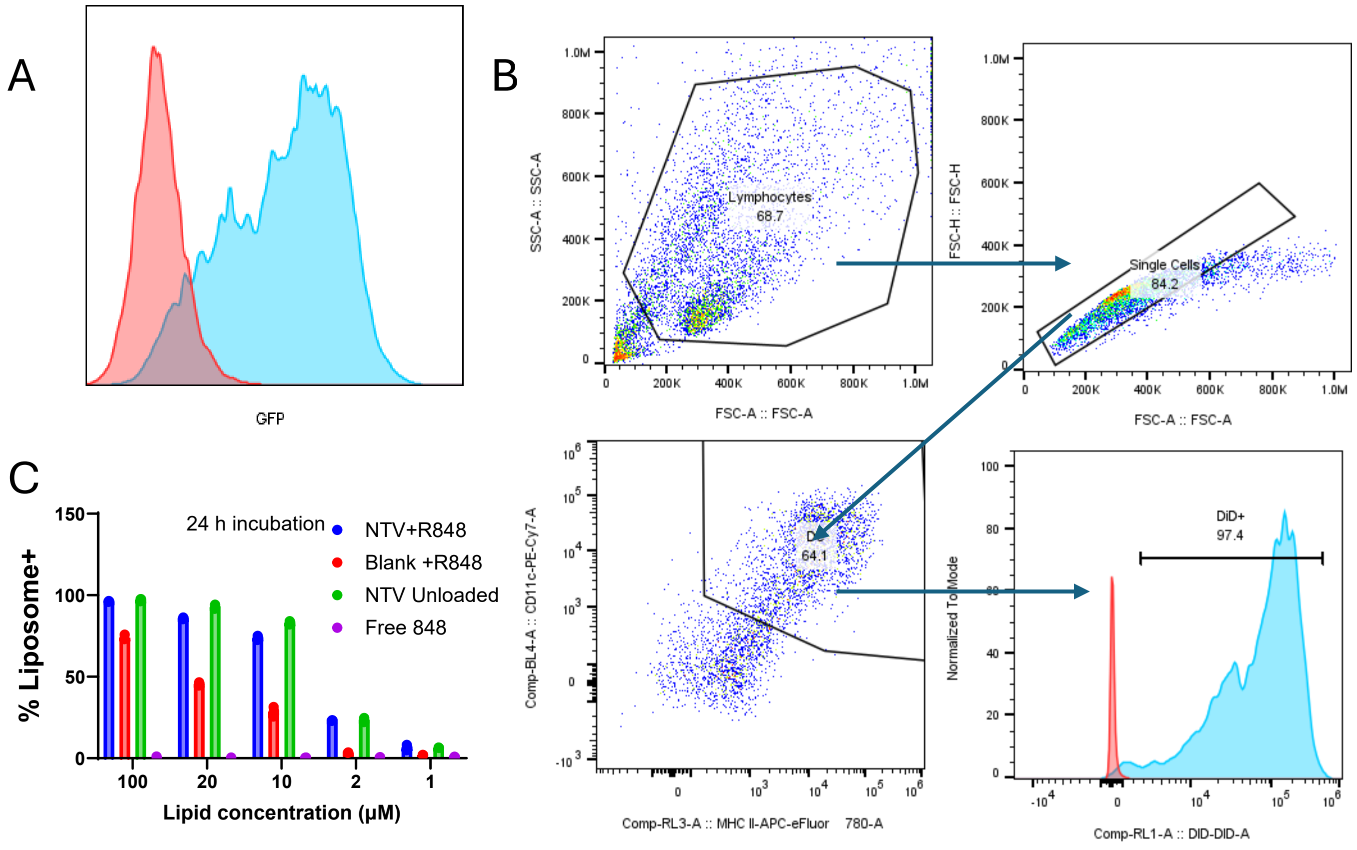


Figure S-12. Additional BMDC uptake data. (A) 100k BMDCs were incubated with 25 ng/mL p24 concentration of unlabeled HXB2 PV (red) or DiO labeled PV for 1 h then analyzed by flow to validate signal. (B) Gating strategy for BMDC experiments in figure 3E/F. (C) As in Figure 3C-F, 10K BMDCs were incubated with NTVs loaded with R848, Blank liposome+ R848, unloaded NTV or free R848. DiD+ cell populations were measured.

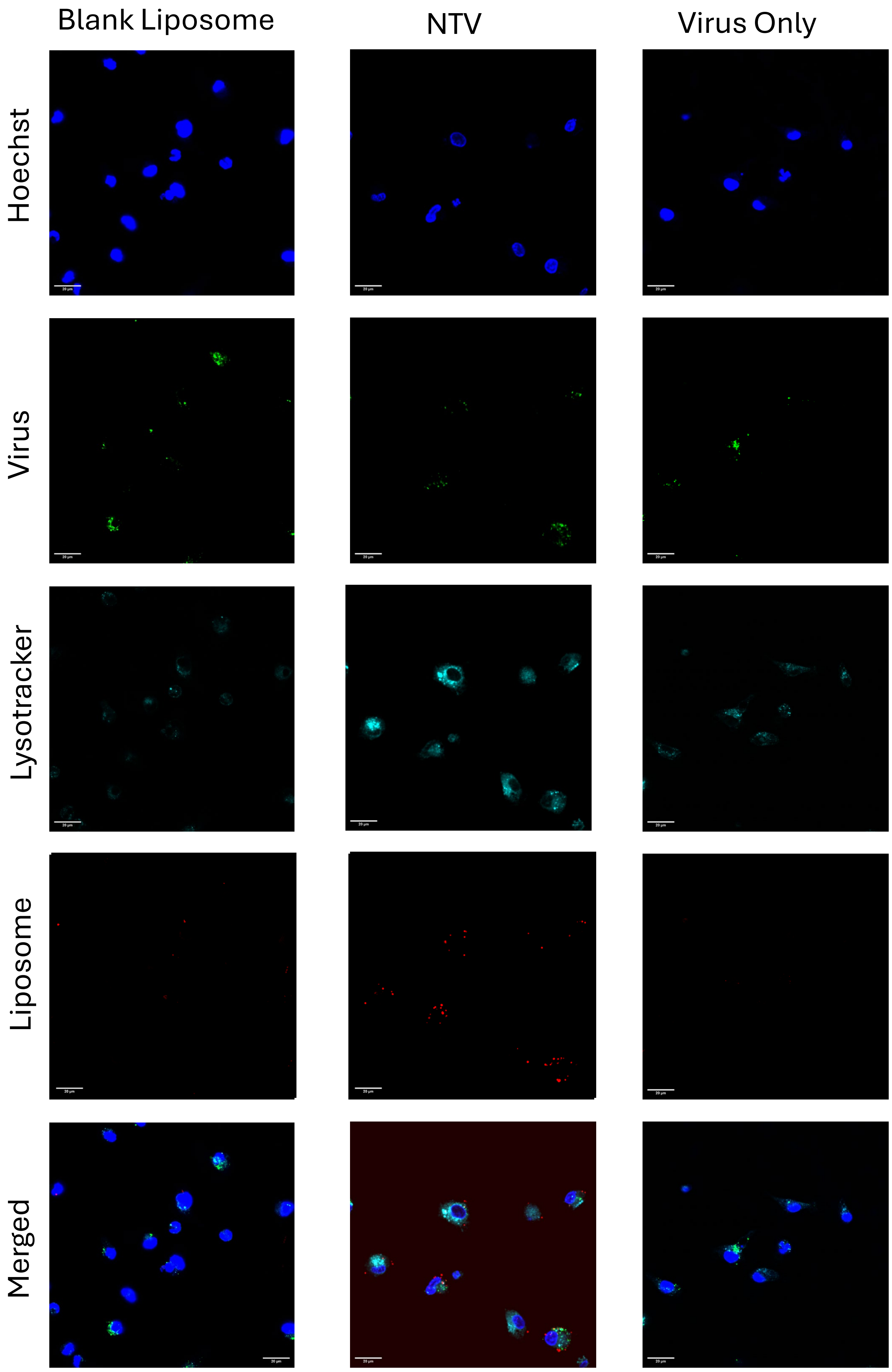


Figure S-13. Representative widefield images of BMDC-Virus-liposome interactions from Figure 3. Scale bars indicate 20 μm.

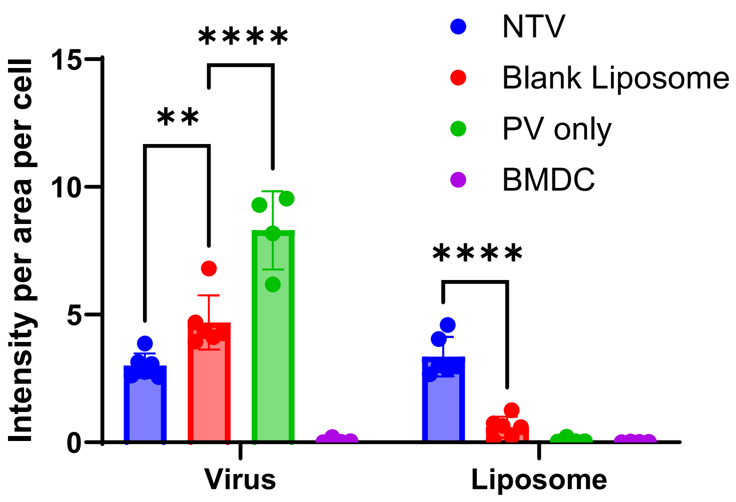


Figure S-14. Average intensity per area per cell from BMDC microscopy images. 4-6 widefield images with 8-20 cells per image were analyzed for average pixel intensity per area and then averaged by number of cells per image for both DiO (virus) signal and DiD (liposome) signal. Error bars ±SD. Significance determined by one way ANOVA. **$p<0.01$, ****$p<1x10^{-5}$

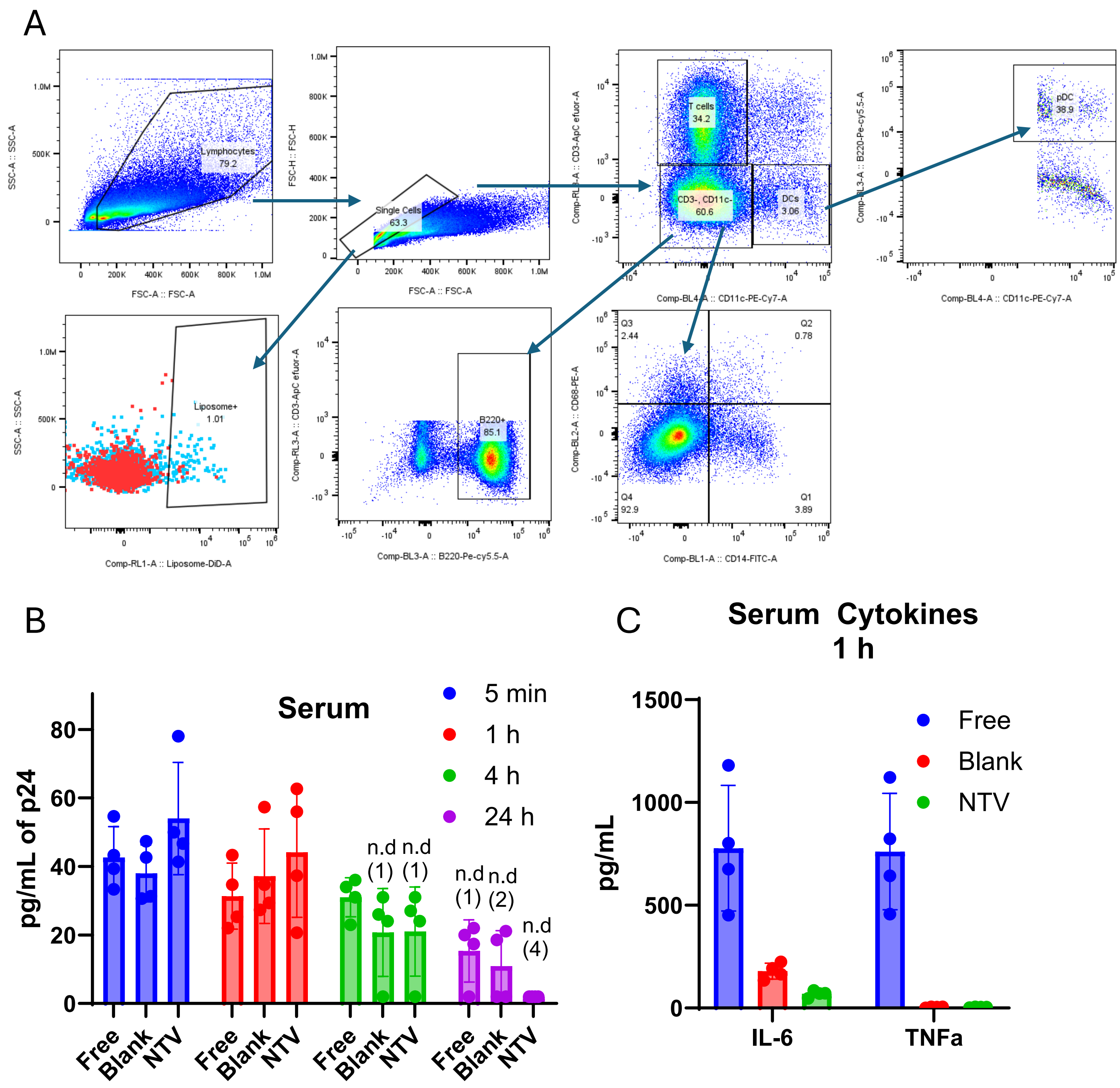


Figure S-15. Additional data for in vivo assay in Fig 4A. (A) Gating Strategy for LN and RBC depleted blood in Fig 4A. Bottom left shows gating for liposome positive (blue) vs FMO control (red). (B) Blood from mice was collected at 5 min, 1 h, 4 h and 24 h after injection of liposome and analyzed by ELISA for p24 antigen. (C) Serum cytokines at 1 h post injection. N=5

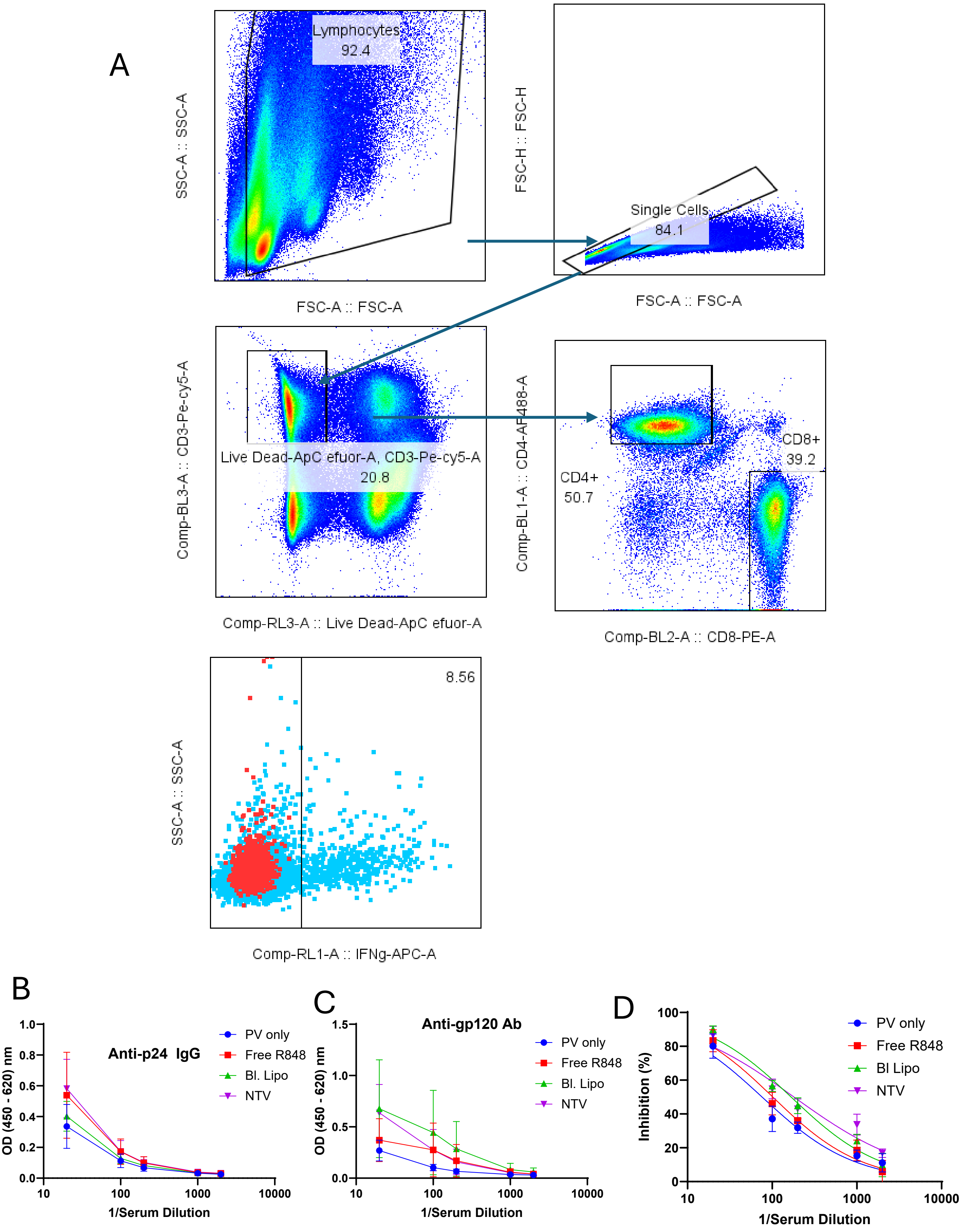


Figure S-16. Additional data for serum analysis of mouse experiment in Figure 4E. (A) Gating strategy. Bottom left is gating for IFNg+ cells (blue) vs FMO control (red). (B) Serum dilutions analyzed by ELISA for p24 or (C) gp120. (D) %GFP+ cells (indicating infection) for GHOST assay with serum dilutions. N=5
